# Recruited macrophages that colonise the post-inflammatory peritoneal niche convert into functionally divergent resident cells

**DOI:** 10.1101/2020.11.30.404988

**Authors:** P. A. Louwe, L. Badiola Gomez, H. Webster, G. Perona-Wright, C. C. Bain, S. J. Forbes, S. J. Jenkins

## Abstract

Inflammation generally leads to substantial recruitment of monocyte-derived macrophages. What regulates the fate of these cells and to what extent they can assume the identity and function of resident macrophages remains unclear. We compared the normal fate of inflammation-elicited macrophages in the peritoneal cavity with their potential under non-inflamed conditions and in the absence of established resident macrophages. Following mild inflammation, elicited macrophages persisted for at least 5 months but failed to fully assume a GATA6^hi^ resident identity due to the presence of enduring resident cells. In contrast, severe inflammation resulted in ablation of resident macrophages and a protracted phase wherein the cavity was incapable of sustaining a resident phenotype, yet ultimately elicited cells acquired a mature GATA6^hi^ identity. Elicited macrophages also exhibited divergent features resulting from inflammation-driven alterations to the peritoneal cavity micro-environment and environment-independent features related to origin and time-of-residency. Critically, one environment-dependent feature of inflammation-elicited macrophages irrespective of severity of inflammation was a failure to produce the chemokine CXCL13, which correlated with a progressive loss in accumulation of peritoneal B1 cells post-inflammation. Hence, rather than being predetermined, the fate of inflammation-elicited peritoneal macrophages appears largely regulated by environment, changes in which result in long-term alteration in function of the peritoneal macrophage compartment post-inflammation.

## Introduction

Inflammation radically alters the composition and function of the tissue macrophage compartment, typically leading to substantial recruitment of monocytes from the blood and activation or even loss of the tissue resident cells^1^. While these early cellular processes have been well characterized across various tissues and models of inflammation, it remains poorly understood how homeostasis within the macrophage compartment is reinstated post inflammation and consequently what long-term effects inflammation may have on tissue macrophage function.

In the steady-state resident macrophages across tissues share expression of core lineage-related genes upon which a tissue-specific transcriptional, epigenetic and functional identity is overlaid^2–5^. These unique molecular identities are largely established upon exposure to tissue-specific environmental signals that in turn drive expression of tissue-specific transcription factors. Tissue resident macrophages also have diverse developmental origins^6,7^ with many tissues containing self-renewing populations largely seeded during embryogenesis and short-lived bone marrow (BM)-derived populations that seemingly inhabit distinct anatomical regions^8,9^. However, in the absence of embryonically-seeded macrophages, circulating monocytes appear able to give rise to long-lived and transcriptionally and functionally normal resident cells^10–14^, suggesting origin may not be a key determinant of macrophage identity per se, but rather tissue-specific anatomically-restricted signals and cell interactions constitute a ‘niche’ that controls macrophage longevity and gene expression. Whilst a small number of seemingly ontogeny-related transcriptional differences may distinguish embryonic and monocyte-derived resident macrophages present within the same ‘niche’^12,14^, limited evidence suggests that even these may be gradually reprogrammed over time^13,15^. Based on these observations, it has been proposed that competition for signals that drive survival of macrophages and expression of tissue-specific transcription factors dictates the balance between incumbent resident macrophages and infiltrating monocytes^16^.

Macrophages in the peritoneal cavity regulate peritoneal B1 cells^15,17^ and provide immune surveillance of the cavity^18^ and neighboring tissues^19^ but they are also implicated in many pathologies, including endometriosis, post-surgical adhesions, pancreatitis and metastatic cancer^18^. The peritoneal cavity contains two populations of resident macrophages: an abundant population of so-called ‘large’ peritoneal macrophages (LPM) that are embryonically-seeded and long-lived, and a rarer population of short-lived MHCII^+^ monocyte-derived cells termed small peritoneal macrophages (SPM)^20^. The transcriptional identity of LPM is dependent upon the transcription factors GATA6^5,21,22^ and CEBPβ^23^ while SPM depend upon IRF4^24^. Expression of GATA6 by LPM is driven primarily by omentum derived retinoic acid at least in part via RXrα^25^. Despite initially having an embryonic origin, the LPM population is gradually replaced by monocyte-derived cells with age, a process that occurs more rapidly in males^20^. Thus, differential rates of replenishment alters the time-of-residency of each macrophage, which leads to differences in phenotype and function of LPM between the sexes and with age^15,20^. Indeed, single cell RNA-sequencing of peritoneal macrophages has revealed LPM in both sexes comprise 3 transcriptionally distinct subsets that, at least in females, appear to represent subsets with different times-of-residency ^15^. However, while tissue resident macrophages in solid organs are envisaged to have a static architectural niche comprising stable cell interactions^16,26^ as delineated in several tissues^27–29^, peritoneal macrophages ‘float’ in a fluidic environment^30^ implying more complex interactions control their maintenance, identity and sub-specialization.

Monocyte-derived macrophages recruited during inflammation typically exhibit distinct transcriptional, functional and phenotypic signatures to resident cells, even in tissue sites where resident cells are ordinarily replenished by monocytes^31,32^. However, it’s unclear whether inflammatory macrophages are fully capable of reprogramming to become resident macrophages and hence if their fate is predominantly regulated by access to appropriate ‘niche’ signals or rather predetermined during their initial differentiation. Macrophages recruited to the alveolar space following influenza infection or bleomycin-induced lung damage persist for many months^33,34^, but retain significant transcriptional differences to enduring resident cells. Notably, established resident macrophages exhibit a relatively poor ability to engraft and reprogram upon adoptive transfer into an ectopic tissue site^4,14^, suggesting that differentiation of macrophages may lead to substantial loss in plasticity. Irrespective, if reprogramming of inflammatory macrophages also has an element of time-dependence, their persistence would be predicted to lead to prolonged alteration in the functional capacity of the tissue macrophage compartment.

In the peritoneal cavity, sterile inflammation can cause substantial contraction in number of LPM through cell death or loss in fibrin clots ^30,35^ but the extent of this loss appears dependent on stimulus and severity of inflammation^35,36^. While remaining LPM can subsequently proliferate during the resolution phase^37^, peritoneal inflammation ^38,39^ including that caused by abdominal surgery^15^ can lead to at least partial replacement of the long-lived LPM population from the BM, with the degree of replacement seeming to correlate with the extent of the preceding loss of incumbent cells^35^. The functional implications of displacement of the resident population remains unclear.

Here we studied the peritoneal cavity, a clinically relevant site commonly used to model inflammatory processes, to investigate what regulates the fate of inflammatory macrophages following sterile inflammation. Using adoptive transfer to unequivocally track inflammatory macrophages and determine the degree to which the environment dictates the fate of these cells, we demonstrate that macrophages infiltrating the cavity following mild inflammation persist long-term but that competition with incumbent resident macrophages inhibits effective acquisition of a mature resident phenotype. Consistent with this competition model, severe inflammation, which caused ablation of incumbent resident macrophages resulted in conversion of inflammatory macrophages to mature resident cells. We therefore reveal the existence of a ‘biochemical niche’ for resident peritoneal macrophages. Competition for the ‘niche’ largely dictates the capacity of monocyte-derived cells to undergo conversion to mature resident macrophages and a failure to compete retains them in a highly proliferative and immunoregulatory state.

## Results

### Inflammatory macrophages persist following mild resolving peritoneal inflammation

To investigate what regulates the fate and phenotype of inflammatory and resident macrophages following resolution of sterile peritoneal inflammation, we used a well-characterised model of intraperitoneal injection with low dose zymosan A (10μg/mouse), in which both populations remain present following resolution of the neutrophilic phase^36,37,40^. First, to definitively delineate incumbent resident cells from inflammatory macrophages recruited during the acute phase of inflammation, we utilized an established method of injecting fluorescent PKH26-PCL dye intraperitoneally 24hrs before zymosan to exclusively label peritoneal phagocytes present prior to inflammation^38^. Uptake of PKH26-PCL dye was largely restricted to all resident LPM and most SPM **(Supplementary Figure 1a)**, identified as F4/80^Hi^ or F4/80^Lo^ CD226^+^ cells respectively^24,40^, and no free dye remained 24hrs later **(Supplementary Figure 1b,c)**. Subsequent injection of low-dose zymosan induced disappearance of dye-labelled (Dye^Hi^) F4/80^Hi^ resident macrophages and influx of dye-negative (Dye^Lo^) Ly6c^+^ monocytes and neutrophils within 4hrs. By day 3, neutrophils were largely cleared and remaining Dye^Lo^ infiltrating cells now exhibited a predominantly F4/80^Int^ Ly6c^Lo/Int^ phenotype consistent with inflammatory macrophages^31,36^ and dye^Hi^ F4/80^Hi^ resident macrophages had partially recovered in number **(Figure 1a,b)**, consistent with their reported repopulation by self-renewal in this model^37^. Finally, to validate our dye-based tracking system, we assessed dye-labelling in tissue-protected BM chimeric mice which allow recruited and resident cells to be determined definitively^20^. This confirmed that Dye^Lo^ F4/80^Int^ macrophages present in the peritoneal cavity at day 3 were derived from recruited cells as evidenced by their high levels of non-host chimerism, whereas Dye^Hi^ F4/80^Hi^ cells displayed low levels of chimerism, demonstrating their tissue residency **(Supplementary Figure 1d)**. Moreover, Dye^Lo^ F4/80^Int^ had high levels of MHCII and virtually no expression of the resident macrophage marker Tim4 **(Supplementary Figure 1e)**, features known to differentiate inflammatory from resident macrophages during resolution^31,36^. Thus, PKH26-PCL-labelling faithfully delineated resident vs recruited macrophage subsets and importantly, this system used a minimal number of surface antibodies thereby circumventing potential confounding effects of adoptive transfer of antibody-coated cells.

**Figure 1.**
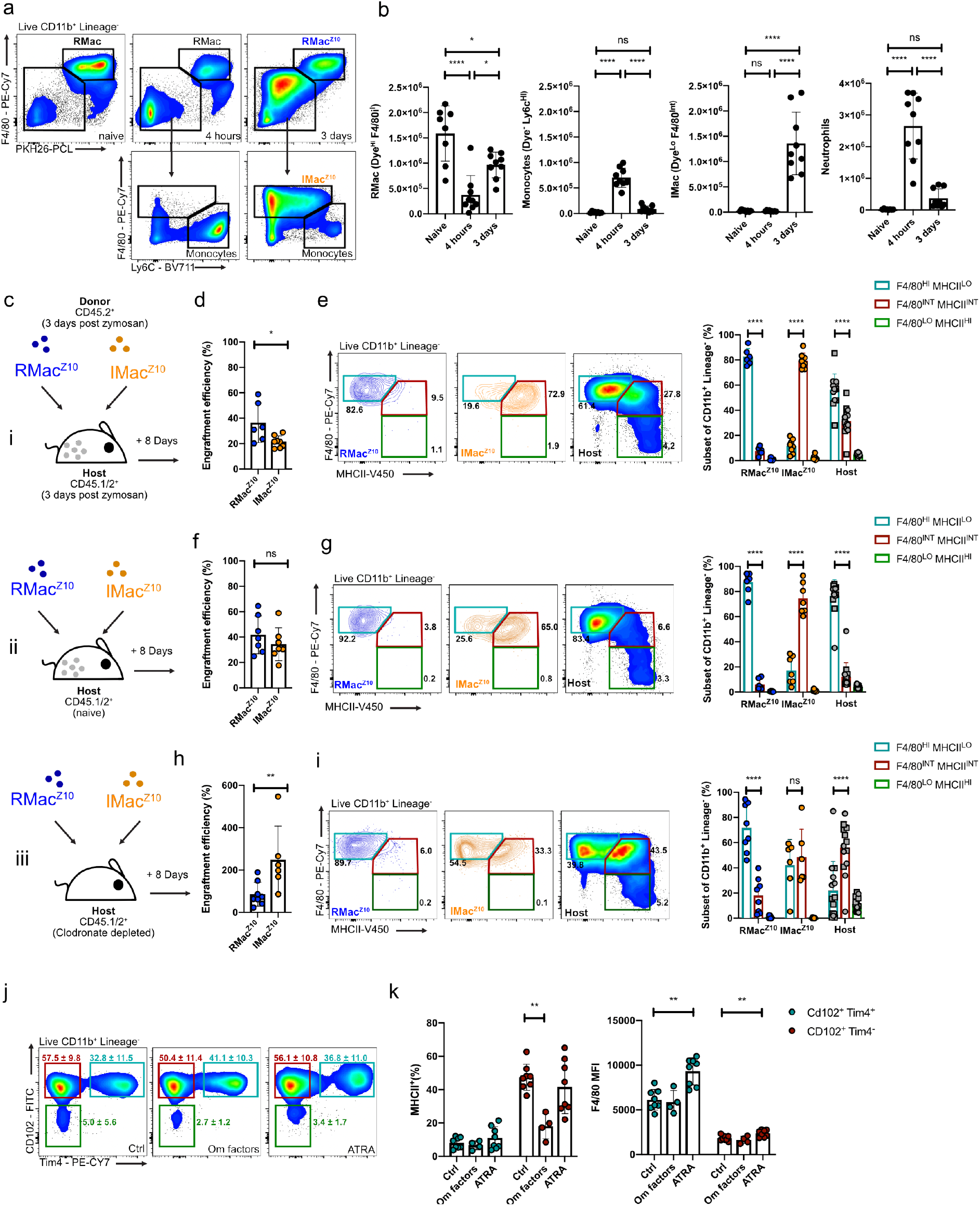
Competition mediates inflammatory macrophage phenotype early post resolution. **(a)** Representative expression of F4/80, Ly6C and PKH26-PCL labelling and identification of F4/80^HI^ PKH26-PCL^HI^ resident macrophages, PKH26-PCL^LO^ Ly6c^+^ monocytes and PKh26-PCL^LO^ F4/80^INT^ inflammatory macrophages in the naïve peritoneal cavity, 4hrs and 3d post 10μg zymosan. **(b)** Absolute numbers of RMac, Monocytes, IMac and Neutrophils the naïve peritoneal cavity (n=8), 4hrs post zymosan (n=9) and 3d post zymosan (n=9). *<0.05, ****p<0.0001 determined by one-way ANOVA with Tukey’s multiple comparisons test. (**c**) Experimental scheme for the adoptive transfer of RMac^Z10^ (blue) or IMac^Z10^ (orange) sourced from CD45.2 mice 3d after injection of 10μg zymosan into mirroring inflamed (i), naïve (ii) or macrophage-depleted (iii) CD45.1/2 recipient mice. (**d**) Engraftment efficiency of transferred RMac^Z10^ (n=6) and IMac^Z10^ (n=8) 8d after transfer into mirroring inflamed recipients. *p<0.05 determined by Mann-Whitney test. **(e)** Expression of F4/80 and MHCII by donor RMac^Z10^ (n=6), IMac^Z10^ (n=8) or host (n=14) cells 8d post transfer. ****p<0.0001 determined by two-way ANOVA and post hoc Tukey test (**f**) Engraftment efficiency of transferred RMac^Z10^ (n=7) and IMac^Z10^ (n=7) 8d after transfer into naïve recipients. **(g)** Expression of F4/80 and MHCII by donor RMac^Z10^ n=7), IMac^Z10^ (n=7) or host (n=14) cells 8d after transfer.****p<0.0001 determined by two-way ANOVA and post hoc Tukey test (**h**) Engraftment efficiency of transferred RMac^Z10^ (n=8) and IMac^Z10^ (n=6) 8d after transfer into clodronate depleted recipients. *p<0.05 determined by Mann-Whitney test. **(i)** Expression of F4/80 and MHCII by donor RMac^Z10^ (n=8), IMac^Z10^ (n=6) or host (n=14) cells 8d after transfer. ****p<0.0001 determined by two-way ANOVA and post hoc Tukey test **(j)** Representative expression of CD102 and Tim4 by cultured cells after 24hrs culture with indicated treatment. **(k)** Proportion of macrophage subsets that express MHCII and F4/80 MFI after 24hrs culture with indicated treatment. **p<0.01 determined by one way Anova and Dunnet’s multiple comparisons test for each subset individually, followed by Bonferroni adjustment. For all experiments, data are presented as mean ± standard deviation with each symbol representing an individual animal, except for (l) where symbols represent individual culture wells. All data were pooled from 3 independent experiments except for (k,l), which were pooled from 2 experiments. Where presented, host cells represented by squares or circles are from animals given RMac^Z10^ or IMac^Z10^ respectively.

Next, we used adoptive transfer to unequivocally determine the fate of these populations. Specifically, Dye^Hi^ F4/80^Hi^ resident macrophages (RMac^Z10^) and Dye^Lo^ F4/80^Int^ inflammatory macrophages (IMac^Z10^) were FACS-purified from C57BL/6 WT (CD45.2^+^) donor mice 3 days after injection of low dose zymosan **(Figure 1a)** and transferred intraperitoneally into separate congenic WT (CD45.1/2^+^) host animals. The recipient mice had been pre-treated 3 days prior with an equivalent dose of zymosan to ensure labelled cells were transferred into a similar environment (**Figure ci)**. Eight days post-transfer, transferred donor RMac^Z10^ and IMac^Z10^ exhibited a similar degree of engraftment, defined as the number retrieved as a proportion of those transferred, although this was somewhat greater for RMac^Z10^ **(Figure 1d)**. Whereas transferred RMac^Z10^ remained predominantly MHCII^Lo^, IMac^Z10^ remained largely MHCII^Hi^ and continued to express marginally less F4/80 such that the two donor populations were identified with relative accuracy using these markers **(Figure 1e)**. Critically, virtually all transferred IMac^Z10^ expressed the LPM-specific transcription factor GATA6 but at markedly lower levels than RMac^Z10^ **(Supplementary Figure 2a,b)**. The host CD11b^+^ myeloid compartment also contained a mixture of F4/80^Int/Hi^ MHCII^Hi^ GATA6^+^ and F4/80^Hi^ MHCII^Lo^ GATA6^+^ macrophages, consistent with persistence of endogenous inflammatory and resident macrophages, but also a minor fraction of GATA6^−^ F4/80^Lo^ MHCII^Hi^ cells **(Figure 1e; Supplementary Figure 2c)** suggestive of newly generated SPM and/or CD11b^+^ DCs. Hence, by combining dye-labelling and adoptive transfer, we developed a robust system to identify and fate map tissue resident and inflammatory macrophages in the context of peritoneal inflammation. Furthermore, our data reveal that the distinct populations of MHCII^−^ and MHCII^+^ peritoneal macrophages present following zymosan-induced peritoneal inflammation^36^ arise from persistence of tissue-resident macrophages established prior to inflammation and monocyte-derived macrophages recruited at the onset of inflammation, respectively.

### Resident macrophages limit initial survival and phenotype of inflammatory macrophages

We next explored what regulates the short-term fate of these cells. First, we transferred RMac^Z10^ and IMac^Z10^ into naïve recipient mice **(Figure 1cii)** to determine whether their survival and phenotype is dictated primarily by the post-inflammation micro-environment. In this homeostatic environment both donor populations persisted equally **(Figure 1f)**, with a level of engraftment akin to that observed for RMac^Z10^ transferred to inflamed mice **(Figure 1d)**. Despite this, IMac^Z10^ remained MHCII^Hi^ **(Figure 1g)** and expressed intermediate levels of GATA6 **(Supplementary Figure 2d),** suggesting this phenotype was not a product of the post-inflammatory micro-environment.

Next to determine if competition with resident macrophages regulates survival and phenotype of IMac^Z10^, we pre-treated recipient mice 7 days prior to transfer with clodronate-loaded liposomes **(Figure 1ciii)**. This regime caused rapid and prolonged loss of recipient F4/80^HI^ LPM, with the cavity being essentially devoid of these cells at the point of adoptive transfer at day 7 **(Supplementary figure 2e)**. In the absence of endogenous resident macrophages, engraftment efficiency of IMac^Z10^ was approximately 250%, indicating these cells have the ability to expand to fill the empty niche **(Figure 1h)**. Furthermore, nearly 50% of IMac^Z10^ adopted a more resident-like MHCII^Lo^ phenotype **(Figure 1i)** but failed to acquire similar levels of GATA6 as RMac^Z10^ within this period **(Supplementary Figure 2f)**. Surprisingly, although RMac^Z10^ also persisted better in the depleted environment, with an engraftment efficiency nearer 100%, they were unable to expand to the same degree as IMac^Z10^ **(Figure 1h)**. Notably, host macrophages also repopulated the cavity during this period, yet they largely exhibited an MHCII^Hi^ phenotype **(Figure 1i)** resembling that of IMac^Z10^ in their native inflamed environment, suggesting these cells likely derive from Ly6C^+^ monocytes recruited to the cavity post-depletion **(Supplementary Figure 2e)**. We also found that irrespective of environment, nearly all IMac^Z10^ expressed the GATA6-independent LPM marker CD102^5^ yet few expressed Tim4 **(Supplementary Figure 2g)**. Together, these data suggest that while IMac^Z10^ persist through the early phases of resolution, their survival and conversion to MHCII^Lo^ cells is largely regulated by the presence of competing resident macrophages.

GATA6 expression by LPM is largely induced by retinoic acid from omental and peritoneal stromal cells whereas the omentum produces additional factors that can drive retinoic acid-independent features of LPM^5,41^. We therefore cultured peritoneal cells collected 11 days post zymosan with all trans retinoic acid (ATRA) or omentum culture supernatant (Om factors) for 24 hours to determine whether MHCII expression by inflammatory macrophages is responsive to retinoic acid or other omental factors. As CD102 and Tim4 did not appear to be altered in any of the *in vivo* experiments we used these to identify resident (CD102^+^/Tim4^+^) and inflammatory CD102^+^Tim4^−^) macrophages post culture. Indeed, post-culture and expression of these surface markers remained unchanged between treatments **(Figure 1j)**. Culture with ATRA led to increased expression of the GATA6 responsive marker F4/80^5^ by CD102^+^/Tim4^+^ resident and CD102^+^Tim4^−^ inflammatory macrophages but not down-regulation of their MHCII expression. In contrast, culture with omental supernatant led to downregulation of MHCII by CD102^+^Tim4^−^ inflammatory macrophages**(Figure 1k)**. Hence, the presence of competing resident macrophages may limit the differentiation of inflammatory macrophages to a MHCII^Lo^ resident phenotype by restricting availability of RA-independent signals in the omental niche.

### Inflammatory macrophages persist long-term but retain cell-intrinsic and environment-dependent transcriptional differences

To investigate whether IMac^Z10^ can persist long-term and fully assimilate into the resident LPM compartment, we continued to track these and the prevailing RMac^Z10^ following transfer into native inflamed cavities versus macrophage-depleted cavities and assessed their phenotype after 8 weeks. Furthermore, to understand whether inflammation changed the behaviour and long-term fate of RMac^Z10^, we included F4/80^Hi^ Dye^Hi^ LPM from naïve donors (RMac) transferred into naïve mice or macrophage-depleted animals for comparison **(Figure 2a).** Notably, only 60% of the transferred RMac^Z10^ retained PKH26-PCL-labelling by this time while some recipient cells had acquired dye **(Supplementary Figure 3a)** confirming the need for adoptive transfer to accurately discriminate these cells. In these experiments, a similar proportion of transferred IMac^Z10^ persisted in their native environment to both RMac^Z10^ and RMac **(Figure 2b,left)**. However, retrospective pooling of all data generated from this time-point throughout our study **(Figure 2b with 4b)** revealed an overall pattern that was similar to day 8, whereby IMac^Z10^ persisted marginally less well than their RMac^Z10^ counterparts **(Supplementary figure 3b)**. Indeed, the overall similarity in survival of donor IMac^Z10^ and RMac^Z10^between day 8 **(Figure 1d)** and week 8 **(Figure 2b)** post-transfer suggests little loss of either population occurred in this time and demonstrates that macrophages elicited by an inflammatory agent become long-lived resident macrophages. Furthermore, the comparable survival of both IMac^Z10^ and RMac^Z10^ to RMac in naïve mice suggests that as early as day 3 post-zymosan injection the homeostatic mechanisms regulating longevity/autonomy of peritoneal macrophages are re-instated. Following transfer into depleted recipients, IMac^Z10^ again expanded significantly in number whereas RMac^Z10^ did not **(Figure 2b, right)**. Likewise, the similarity in persistence of engrafted IMac^Z10^ and RMac^Z10^ between day 8 **(Figure 1h)** and week 8 post-transfer **(Figure 2b)** suggests that the resident peritoneal macrophage pool also quickly re-establishes following depletion and resumes self-maintenance irrespective of origin.

**Figure 2.**
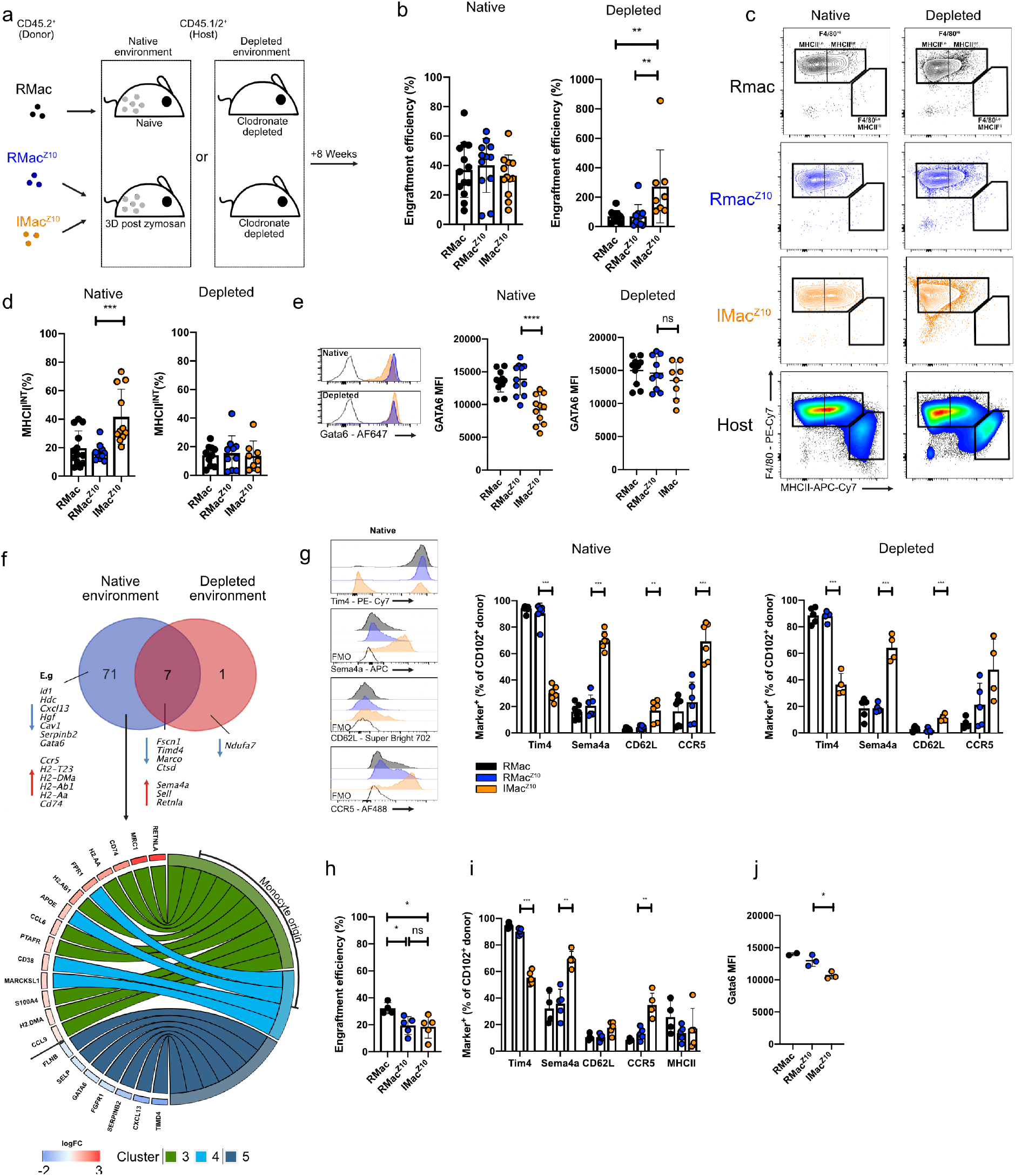
Colonizing inflammatory macrophages are long lived but retain intrinsic and environment-dependent differences to RMac. (**a**) Experimental scheme for the adoptive transfer of RMac from naïve mice or RMac^Z10^ and IMac^Z10^obtained 3d after injection of 10μg zymosan into mirroring naïve, inflamed or clodronate depleted recipients. **(b)** Engraftment efficiency of transferred RMac, RMac^Z10^and IMac^Z10^ 8wk after transfer into the mirroring recipients (left; n=13, n=9, n=9) or depleted recipients (right; n=10, n=10, n=8) **p<0.01 determined by one-way ANOVA and Tukey’s multiple comparisons test **(c)** Representative expression of F4/80 and MHCII of indicated donor populations 8wk after transfer into native (left) or clodronate depleted (right) recipients. Bottom, representative expression of host myeloid cells post zymosan or post depletion. **(d)** Proportion of donor RMac, RMac^Z10^and IMac^Z10^ that express MHCII 8wk after transfer into mirroring recipients (left; n=13,11,11) or depleted recipients (right; 10, 10, 8). **p<0.01 determined by one-way ANOVA and Tukey’s multiple comparisons test **(e)** Mean fluorescence intensity of GATA6 after transfer of RMac, RMac^Z10^ or IMac^Z10^ 8wk after into native (left; n=12,11,11) or depleted (right; n=10,10,8) recipients. ****p<0.0001 determined by one-way ANOVA and Tukey’s multiple comparisons test **(f)** Venn diagram indicating overlap between genes differentially (adj p value <0.05) expressed between RMac^Z10^and IMac^Z10^ 8wk after transfer into the native (blue) or depleted (red) environment. Below, circus plot depicting fold change of differentially expressed genes (left side) that are cluster markers for single cell clusters (right side) identified by Bain et al^15^. **(g)** Expression of markers of interest by CD102^+^ RMac (black), RMac^Z10^ (blue) and IMac^Z10^(orange) 8wk after transfer into mirroring (left; n=7,6,6) or depleted recipients (right; n= 5, 5, 4). **p<0.01, **p<0.01 ***p<0.001 determined by one way ANOVA and Dunnet’s multiple comparisons test for each marker individually, followed by Bonferroni adjustment. **(h)** Engraftment efficiency of transferred RMac, RMac^Z10^ and IMac^Z10^ 22wks after transfer into the mirroring recipients (n=4, 5, 5). *p<0.05 determined by one-way ANOVA and Tukey’s multiple comparisons test **(i)** Expression of markers of interest by CD102^+^ RMac (black), RMac^Z10^ (blue) and IMac^Z10^ (orange) 22wks after transfer into mirroring (n=4, 5, 5) recipients. **p<0.01 ***p<0.001 determined by one way ANOVA and Dunnet’s multiple comparisons test for each marker individually, followed by Bonferroni adjustment. **(j)** Mean fluorescence intensity of GATA6, 22wks after transfer of RMac, RMac^Z10^or IMac^Z10^ into native (n=4, 5, 5) recipients. *p<0.05 determined by one-way ANOVA and Tukey’s multiple comparisons test For all experiments, data are presented as mean ± standard deviation with each symbol representing an individual animal. All data were pooled from at least 2 independent experiments, except for **(j)** which is a single representative experiment. Where presented, host cells represented by squares or circles are from animals given RMac^Z10^or IMac^Z10^ respectively.

Importantly, even after 8wks in their native environment IMac^Z10^ exhibited lower expression of GATA6, marginally less F4/80, and a higher proportion of MHCII^+^ cells than either RMac population **(Figure 2d,e & Supplementary Figure 3d)**. In contrast, prior removal of competing endogenous cells through administration of clodronate liposomes allowed transferred IMac^Z10^ to fully acquire the MHC^Lo^GATA6^Hi^ phenotype of RMac^Z10^ within 8 weeks **(Figure 2d,e & Supplementary Figure 3d)**. To investigate the wider transcriptional integration of IMac^Z10^within the resident LPM compartment, we sorted the 3 donor populations from both native and depleted environments at 8wks post-transfer and investigated gene expression using the nanoString nCounter mouse myeloid panel. This analysis revealed that in their native environment IMac^Z10^ remained highly transcriptionally distinct from RMac^Z10^, with 78 of the 372 detected genes being differentially expressed **(Supplementary Table 1, Figure 2f, Supplementary Figure 3e)** whereas no detectable differences were apparent between RMac and RMAC^10^**(Supplementary Table 2)**. Of the 13 genes included in the panel considered to differentiate LPM from other tissue resident macrophages^5,41^, 11 were differentially expressed between IMac^Z10^ and RMac^Z10^ **(Supplementary Figure 3f)** including *Gata6*. In contrast, *Cebpb*, which encodes the transcription factor CEBPβupon which LPM are also dependent^23^, was expressed equally by IMac **(Supplementary Table 1)**. A quarter of the genes differentially-expressed between IMac^Z10^ and RMac^Z10^ overlapped with those regulated by GATA6 in LPM^5,21,22^, including *Adgre1,* which encodes F4/80, and consequently the gene signature of IMac^Z10^ largely aligned with that of *Gata6*-deficient LPM^5,21,22^ **(Supplementary Figure 3g)**. Furthermore, comparison with our published single cell RNAseq analysis^15^ of LPM revealed that genes expressed more highly in IMac^Z10^ overlapped exclusively with those expressed more highly by LPM of recent monocyte origin in naïve female mice (e.g. *Apoe*, *Retnla*, and genes related to MHCII presentation), whereas genes expressed more highly in RMac^Z10^ overlapped exclusively with those expressed more highly by the most long-lived LPM (e.g. *Timd4*, *Cxcl13*, and *Gata6*) **(Figure 2f)**. Moreover, re-analysis of our single cell RNA-seq dataset of LPM^15^ revealed that cluster markers that define monocyte-derived LPM overlapped markedly and exclusively with genes expressed more highly by *Gata6*-deficient macrophages^5,21,22^ whereas cluster markers of established LPM overlapped substantially and exclusively with genes expressed more highly by *Gata6*-sufficent LPM **(Supplementary Figure 3h)**. Hence, these data suggest that differences in the degree of GATA6 expression between established resident macrophages and incoming monocyte-derived macrophages controls a significant proportion of the genes differentially-expressed between these populations in steady-state and post-inflammation.

Critically, gene expression profiling suggested that IMac^Z10^ and RMac^Z10^ became transcriptionally more similar after transfer into depleted recipients, with the number of differentially expressed genes decreasing from 78 to 8 (**Figure 2f, Supplementary Table 3)**. Notably, differences in *Gata6* and almost all the potentially GATA6-regulated genes were lost **(Supplementary Figure 3i),** as were differences in most other genes that defined clusters identified in steady state^15^, including *Cxcl13* and MHCII-related genes. Unsurprisingly, RMac and RMac^Z10^ remained transcriptionally indistinct in the macrophage-deplete environment (**Supplementary Table 4**). Hence, these data suggest the majority of transcriptional differences between IMac^Z10^ and RMac^Z10^ are determined by the post-inflammatory environment or competition with incumbent resident macrophages for access to niche signals, whereas a smaller number of differentially expressed genes may represent cell-intrinsic features related to origin. Specifically, enduring resident macrophages seemingly prevent IMac^Z10^ transition to a mature GATA6^hi^ phenotype, thus retaining them in a transcriptional state associated with steady-state monocyte-derived LPM.

Flow-cytometric analysis confirmed that within the native post-inflammatory environment, IMac^Z10^ expressed higher levels of Sema4a, CD62L, and CCR5 and but largely failed to acquire expression of Tim4 **(Figure 2g, left)**, consistent with the differential expression of *Sema4a*, *Sell* (encoding for CD62L), *Ccr5*, and *Timd4* detected by nanoString. Similarly, IMac^Z10^ in the depleted environment retained equivalently high levels of Sema4a and CD62L, and low levels of Tim4 expression **(Figure 2g, right)**, consistent with these being cell-intrinsic rather than environment-dependent features of IMac^Z10^ **(Figure 2f)**. In line with an expression pattern predominantly dictated by environmental cues, we found that IMac^Z10^ expressed more variable and on the whole lower levels of CCR5 in the macrophage-depleted environment **(Figure 2g, right)**. We extended this analysis to include surface markers that define newly monocyte-derived (Folate receptor β (FRβ)) and long-lived resident peritoneal macrophages (CD209b and V-set immunoglobulin domain-containing 4 (VSIG4))^15^. IMac^Z10^ failed to acquire equivalent expression of CD209b or VSIG4 to either RMac^Z10^ population irrespective of environment, whereas they exhibited comparatively high levels of FRβ in the native environment that, like CCR5, was lost in macrophage-depleted recipients suggesting down-regulation by environmental cues **(Supplementary Figure 3j)**.

Finally, we determined whether reprogramming of IMac^Z10^ may occur naturally over the lifespan following inflammation. Fate-mapping for 5mths revealed continued persistence of IMac^Z10^, RMac^Z10^ and RMac transferred into their native environments **(Figure 2h),** although only RMac appeared to survive as well as at 8wks **(Figure2b)**. Within this time IMac^Z10^ had downregulated MHCII to levels equivalent to RMac^Z10^, yet they continued to express lower levels of GATA6 and retain higher proportions of cells expressing Sema4a, CD62L, CCR5 **(Figure 2i,j)** and FRβ **(Supplementary Figure 3k)**. Despite this, fewer IMac^Z10^ expressed CCR5 or FRβ than at 8wks **(Figure 2g ‘native’ vs 2i and Supplementary Figure 3j ‘native’ vs 3k)**, consistent with gradual reprogramming of expression of these markers, whereas expression of Sema4a and CD62L remained unchanged. Furthermore, IMac^Z10^ had acquired equivalent levels of VSIG4 to RMac^Z10^ by this time but failed to upregulate expression of Tim4 and CD209b to levels observed on the resident populations **(Figure 2i, Supplementary Figure 3k)**, despite the frequency of IMac^Z10^ expressing these markers increasing compared to 8wks **(Figure 2i to Supplementary Figure 3k).** Of note, the low frequency of RMac and RMac^Z10^ that expressed CCR5 and FRβ by week 8 of transfer was seemingly reduced even further by 5mths, while proportion that expressed Vsig4 and CD209b continued to rise gradually **(Figure 2g ‘native’ vs 2i; Supplementary Figure 3ij’naïve’ vs 3k)**. These data are consistent with our previous supposition that expression of Tim4, CD209b and VSIG4 by LPM is regulated by time-of-residency and demonstrate this remains so following mild inflammation. Hence, the distinct phenotype of IMac-derived LPM appear to comprise: 1) pre-determined features seemingly retained over time and not reprogrammed by niche signals (CD62L, Sema4a); 2) features that fail to reprogram due to an inability to compete with RMac^Z10^ for environmental cues but that are reprogrammed with time (MHCII, GATA6, CCR5, FRβ); and 3) features related to time-of-residency irrespective of competition with RMac^Z10^ (VSIG4, Tim4, CD209b).

### Colonizing Inflammatory macrophages are functionally distinct and resemble monocyte derived resident macrophages

To determine whether colonizing inflammatory macrophages differ functionally and behaviourally to established resident macrophages, we developed a gating strategy based on a Tim4^+^ Sema4a^−^ (R1) and Tim4^−^ Sema4a^+^ (R3) profile to identify the majority of RMac^Z10^ and IMac^Z10^, respectively **(Figure 3a,b).** Using this approach, we were able to track the major long-term changes in phenotype of the resident LPM pool triggered by inflammation without need for dye-based fate-mapping **(Supplementary Figure 4b)**. In addition, to determine whether IMac-derived LPM are functionally similar to LPM of recent monocyte origin recruited during homeostasis, we confirmed that the Tim4^−^Sema4a^+^ gate identified the majority of Tim4^−^ MHCII^+^ LPM in naïve mice (**Supplementary Figure 4c)**, which we previously validated to identify newly monocyte-derived LPM^15^.

**Figure 3.**
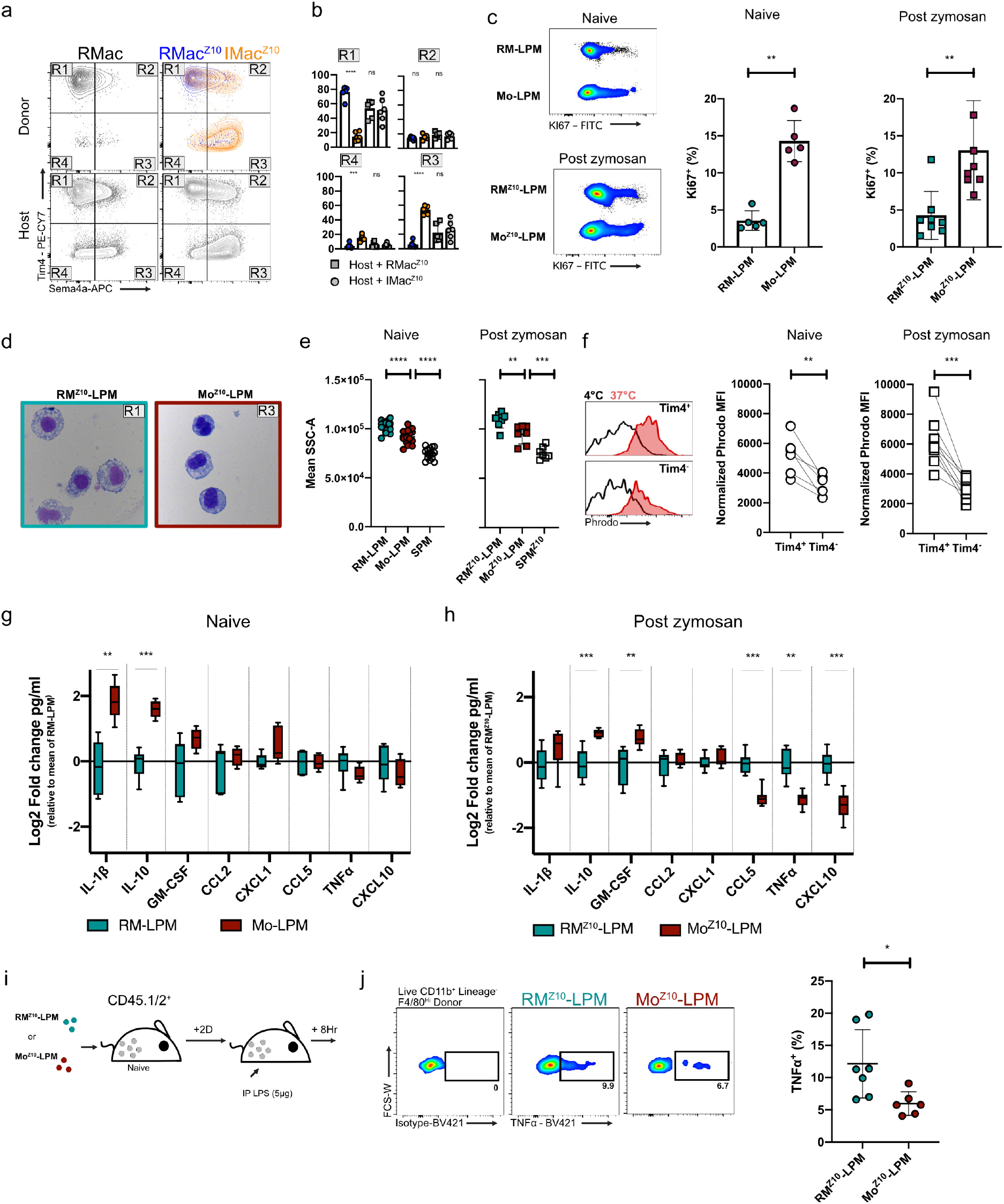
Monocyte derived LPM are functionally distinct from embryonically seeded LPM. **(a)** Representative expression of Tim4 and Sema4a by donor RMac, RMac^Z10^and IMac^Z10^ 8wks post transfer and their respective hosts (bottom) **(b)** Proportion of donor RMac^Z10^ (blue circle, n=6), IMac^Z10^ (orange circle, n=6) and their respective host (grey) macrophages with Sema4a^LO^Tim4^+^(R1), Sema4a^HI^Tim4^+^(R2), Sema4a^Hi^Tim4^−^ (R3) or Sema4a^Lo^Tim4^−^ (R4) phenotype.. ***p<0.001, ****p<0.0001 determined by one way ANOVA and Tukey’s multiple comparisons test **(c)** Expression of Ki67 by naïve RM-LPM and Mo-LPM (n=5) or 8wk post zymosan RM^Z10^-LPM and Mo^Z10^-LPM (n=8). *p<0.05,**p<0.01 ****p<0.0001 determined by one way ANOVA and Tukey’s multiple comparisons test **(d)** Morphological appearance of RM^Z10^-LPM and Mo^Z10^-LPM purified 8wks post zymosan. **(e)** Mean side scatter of naïve RM-LPM, Mo-LPM, SPM (n=5) or 8wks post zymosan RM^Z10^-LPM, Mo^Z10^-LPM and SPM^Z10^(n=8). **p<0.01,***p<0.001 ****p<0.0001 determined by one way ANOVA and Tukey’s multiple comparisons test **(f)** Uptake of Phrodo E.coli particles by naïve (n=6) or 8wks post zymosan (n=9) Tim4^+^ and Tim4^−^ macrophages shown as normalized Phorodo mean fluorescence intensity (MFI 37°C-MFI 4°C). **p<0.01,***p<0.001 determined by paired student’s t test. **(g)** Secreted cytokine/chemokine profile collected from cultures of RM-LPM (n=6, teal) or Mo-LPM (n=5, red) sourced from naïve animals 14hrs after culture with LPS (1ng/ml). Results are shown as log2 fold change in mean pg/ml over the mean RM-LPM using a box-and-whiskers plot. **p<0.001***p<0.0001 determined by repeated student’s t test with Holm-Sidak correction. **(h)** Secreted cytokine/chemokine profile collected from cultures of RM^Z10^-LPM (n=8, teal) or Mo^Z10^-LPM (n=8, red) sourced 8wks post zymosan, 14hrs after culture with LPS (1ng/ml). Results are shown as log2 fold change in mean pg/ml over the mean RM^Z10^-LPM using a box-and-whiskers plot. **p<0.001***p<0.0001 determined by repeated student’s t test with Holm-Sidak correction. **(i)** Experimental scheme for the adoptive transfer of RM^Z10^-LPM and Mo^Z10^-LPM purified from donor mice 8 weeks post 10μg zymosan into naïve recipient mice followed by intraperitoneal treatment with 5μg LPS. **(j)** Proportion of donor CD45.2^+^ F4/80^Hi^ RM^Z10^-LPM(n=7) and Mo^Z10^-LPM (n=6) 8 hours post LPS that are TNFα positive. *p<0.05 determined by student’s t test. For all experiments, data are presented as mean ± standard deviation with each symbol representing an individual animal. Naïve animals were age matched to zymosan treated (15-18Wk) animals except for (c,g) where naïve animals were 10-12Wk at the time of analysis. All data were pooled from 2 independent experiments.

Consistent with our previous observations showing that LPM of recent monocyte-origin proliferate more than established LPM during homeostasis^20^, the Sema4a^+^Tim4^−^ fraction of LPM from naïve mice (subsequently referred to as **Mo-LPM** and **RM-LPM** respectively) exhibited the highest level of proliferation, as determined by Ki67 expression **(Figure 3c)**. Similarly, Sema4a^+^ Tim4^−^ and Sema4^−^ Tim4^+^ defined-populations found 8wks post-zymosan injection (subsequently referred to as **Mo^Z10^-LPM** and **RM^Z10^-LPM** respectively) exhibited the same divergent pattern in proliferative activity (**Figure 3c**). Furthermore, while both Mo^Z10^-LPM and RM^Z10^-LPM displayed typical macrophage morphology, the cytoplasm of RM^Z10^-LPM contained many more vacuoles **(Figure 3d)** indicative of greater phagocytic activity. Indeed, both Mo^Z10^-LPM and Mo-LPM had appreciably lower side-scatter characteristics than their RM counterparts, albeit higher than SPM **(Figure 3e)**. Moreover, examination of phagocytic potential *in vitro* using pHrodo-labelled *Escherichia coli* particles revealed that Tim4^+^ LPM from naïve mice and 8wks after inflammation were significantly more phagocytic than the Tim4^−^ fraction **(Figure 3f)**. Of note, incubation at 37°C for 1hr caused rapid acquisition of surface Sema4a by Tim4^+^ macrophages thereby preventing analysis of Sema4a-defined populations in this assay. Furthermore, re-analysis of our short-term transfer experiments revealed that only Tim4^+^ recipient LPM acquired PKH26-PCL dye from donor RMac irrespective of whether recipients were naïve or zymosan-injected **(Supplementary Figure 4d,e)** suggesting up-take of dying donor cells is restricted to Tim4^+^ cells. Lastly, to test responsiveness to challenge, RM-LPM and Mo-LPM from naïve mice and RM^Z10^-LPM and Mo^Z10^-LPM were purified 8wks post-zymosan injection, exposed *in vitro* to LPS and cytokine and chemokine production assessed by multiplex assay. The overall response of Mo^Z10^-LPM and Mo-LPM compared to their RM counterparts was remarkably similar; both produced higher levels of IL-10 and somewhat more IL-1βand GM-CSF and less CXCL10 and TNFα **(Figure 3g,h)**, suggesting these are common features of monocyte-derived LPM. Furthermore, direct comparison confirmed that despite some subtle differences, Mo-LPM and Mo^Z10^-LPM produced largely similar quantities of cytokines and chemokines, as did RM-LPM compared with RM^Z10^-LPM **(Supplementary Figure 4f,g)**. Hence, together with our gene expression profiling, these data suggest that recency-of-monocyte origin more strongly influences the behaviour of LPM than prior experience of inflammation and that persistence of inflammatory macrophages leads to the expansion of a normally minor subset of IL-10 producing monocyte-derived LPM present under homeostatic conditions. Finally, we found that purified Mo^Z10^-LPM transferred into naïve recipient mice that subsequently received LPS produced less TNFα than transferred RM^Z10^-LPM **(Figure 3i,j)**, confirming these cells also respond differently to challenge *in vivo.*

### Fate of inflammatory macrophages is dependent on the severity of inflammation

In the mild model of sterile peritonitis studied so far, the initial macrophage ‘disappearance reaction’ and inflammatory response that occurs is relatively limited and transient^36^. In contrast, injection of a 100-fold higher dose of zymosan (1000ug/mouse) induced an almost complete and protracted disappearance of F4/80^hi^ Tim4^+^ LPM concurrent with a greater and more protracted influx of monocytes and neutrophils^42^ and overall increase in size of the CD11b^+^ macrophage/monocyte compartment **(Supplementary Figure 5a)**. Notably, the CD11b^+^ population remained exclusively F4/80^lo^ MHCII^Hi^ for at least 11 days although Tim4^+^ cells had begun to re-emerge within this time **(Supplementary Figure 5b)**. Importantly, analysis in tissue-protected BM chimeric mice confirmed that the entire peritoneal macrophage pool, including Tim4^+^ cells, had been replaced from the BM 3wks after high-dose zymosan **(Figure 4a, Supplementary Figure 5c)**. Thus, severe sterile peritoneal inflammation is a physiological setting leading to the complete ablation and replacement of resident LPM.

**Figure 4.**
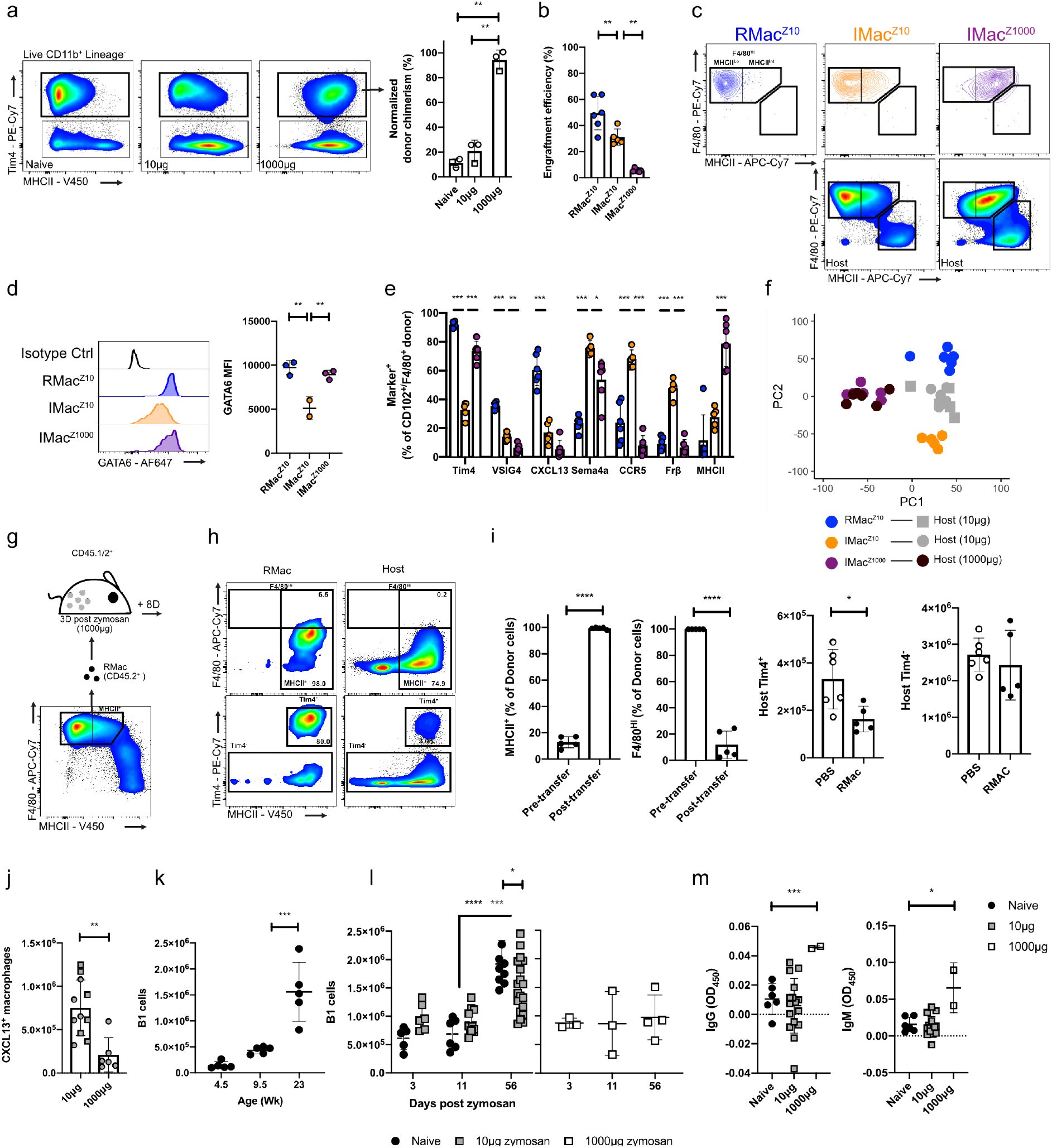
Ontogeny does not dictate monocyte phenotype after severe peritonitis in females and leads to impaired B1 cell expansion. **(a)** Non host chimerism of Tim4^+^ macrophages 17d after indicated zymosan dose in tissue-protected BM chimeric mice. Zymosan treatment 8 (circle) or 26 (square) wks after irradiation. **p<0.01 determined by one-way ANOVA and Tukey’s multiple comparisons test **(b)** Engraftment efficiency of transferred RMac^Z10^ (n=6), IMac^Z10^ (n=5), and IMac^Z1000^ (n=6),after transfer into the mirroring recipients. **p<0.01 determined by one-way ANOVA and Tukey’s multiple comparisons test **(c)** Representative expression of F4/80 and MHCII of indicated donor populations 8wks post after transfer. Bottom, representative expression of host myeloid cells post zymosan or post depletion. **(d)** Mean fluorescence intensity of GATA6 of donor RMac^Z10^, IMac^Z10^ and IMac^Z10^ after transfer into native recipients (n=3,2,3). **p<0.01 determined by one-way ANOVA and Tukey’s multiple comparisons test **(e)** Expression of markers of interest by CD102^+^ or F4/80^+^ donor RMac^Z10^ (blue), IMac^Z10^ (orange) and IMac^Z1000^ (purple) 8wks after transfer into mirroring (left; n=6,5,6) recipients. *p<0.05 **p<0.01 ***p<0.001 determined by one way ANOVA and Dunnet’s multiple comparisons test for each marker individually, followed by Bonferroni adjustment. **(f)** Principal component analysis based on all markers assessed in **(e)** on indicated cell populations. **(g)** Experimental scheme for adoptive transfer of F4/80^Hi^ MHCII^Lo^ naïve resident macrophages (RMac) **(h)** Representative expression of F4/80 (top),Tim4 (bottom) and MHCII on transferred RMac and host myeloid cells 8d post transfer. **(i)** Proportion of RMac that express MHCII and F4/80 (set to 100% pre-transfer as cells were sorted on this marker) before transfer and 8d post-transfer (n=5; left) and absolute number of host Tim4^+^ or Tim4^−^ macrophages 8d post-transfer (n=5). *p<0.05 **\*\*\*\*** p<0.0001determined by student’s t test. **(j)** Absolute number of host macrophages that express CXCL13 8wks after 10μg of zymosan (n=11) or 1000μg zymosan (n=6). **p<0.01 determined by student’s t test. **(k)** Absolute number of peritoneal B1 cells in naïve female mice at indicated age in weeks (n=5/timepoint). ***p<0.001 determined by one-way ANOVA and post hoc Tukey test **(l)** Absolute number of peritoneal B1 cells at indicated timepoints in naïve (black circle; n= 5,6,10), post 10μg zymosan (grey square; n=6,11,20) and post 1000μg zymosan (white square; n=3,3,4) *p<0.05, ***p<0.001, ****p<0.0001 determined by two-way ANOVA and post hoc Tukey test. **(m)** Detection of serum anti phosphorylcholine specific IgM and IgG using enzyme linked immunosorbent assay from naïve (n=6), 8wks post low dose zymosan (n=16) and 8wks post high dose zymosan (n=2). *p<0.05 ***p<0.001 determined by one way ANOVA and Dunnet’s multiple comparisons test For all experiments, data are presented as mean ± standard deviation with each symbol representing an individual animal. All data were pooled from at least 2 independent experiments except high dose presented in **(l,m)** which is from a single experiment. Where presented, host cells represented by squares or circles are from animals given RMac^Z10^ or IMac^Z10^ respectively.

Hence, to understand the fate of inflammatory macrophages after severe inflammation we purified dye-negative inflammatory macrophages 3 days after injection of high or low-dose zymosan **(IMac^Z1000^; Supplementary Figure 5d)** and transferred them into their native inflammatory environments. Markedly fewer donor IMac persisted at 8wks in recipients of high dose zymosan compared with those receiving low dose zymosan **(Figure 4b)**, consistent with the greater contraction in size of the peritoneal macrophage compartment **(Supplementary Figure 5a)** and the reported death of the majority of inflammation-elicited macrophages that follows resolution of severe peritoneal inflammation^31,43^. However, those surviving IMac^Z1000^ in the high dose environment adopted a F4/80^Hi^GATA6^+^ profile by 8wks following severe inflammation and almost none subsequently expressed CCR5 or FRβ**(Figure 4c,d,e)**. These results are consistent with a more mature phenotype, again reflecting more rapid differentiation in the absence of competition from enduring resident macrophages. Nevertheless, a shared deficiency of IMac-derived cells in both high and low-dose environments was the failure to produce CXCL13, a GATA6^21,22^ and Rxra^25^ independent feature of LPMs. Thus, these data suggest impaired CXCL13 expression by IMac arises from long-term alterations in the LPM niche that occurs irrespective of inflammation severity and retinoic acid production. IMac^Z1000^ and IMac^Z10^ derived cells also largely shared the propensity to express the intrinsic marker of monocyte-derived LPM Sema4a **(Figure 4e)**, and to lack expression of the environment-independent but time-dependent marker VSIG4 **(Figure 4e)**. More surprisingly, IMac^Z1000^ retained high levels of MHCII, and largely expressed the otherwise time-dependent marker Tim4 **(Figure 4e)**. Furthermore, donor-derived macrophages following severe inflammation almost perfectly resembled the phenotype of host cells, consistent with their likely uniform origin from inflammatory macrophages **(Supplementary Figure 5e)**. In contrast, in the lower dose environment host macrophages neither aligned with RMac^Z10^ nor IMac^Z10^ but corresponded to a mixed population of these cells, **(Figure 4f, Supplementary Figure 5e)** re-emphasising that phenotype is ontogeny-restricted in this environment. Consequently, the LPM compartment on the whole 8 weeks after high dose zymosan differed markedly to that after low dose zymosan for each marker assessed **(Supplementary Figure 5e)**.

As both F4/80 and MHCII expression by IMac^Z10^ were rapidly responsive to niche signals and competition with LPM after low dose zymosan **(Figure 1k, Figure 2d)**, we postulated that the initially prolonged absence of F4/80^hi^ LPM, rapid acquisition of Tim4 expression, and persistent expression of MHCII by recruited cells **(Supplementary Figure 5b)** following severe inflammation arose from an altered cavity environment. To test this, we adoptively transferred 4×10^5^ F4/80^Hi^, largely MHCII^Lo^, resident macrophages from naïve mice (RMac) into recipient mice 3 days after injection of high dose zymosan **(Figure 4g)**. Eight days later transferred cells had almost exclusively upregulated MHCII expression and markedly down-regulated expression of F4/80 **(Figure 4h,i).** Moreover, transfer of RMac suppressed the rapid acquisition of Tim4 by host cells, as indicated by a specific decrease in number of Tim4^+^ host macrophages **(Figure 4i)**. Hence, novel environmental cues following severe inflammation directly drive expression of MHCII, and in conjunction with the absence of embryonically seeded Tim4^+^ resident macrophages allow rapid acquisition of Tim4 by monocyte-derived cells. Furthermore, there appears to be a phase of at least 11 days during severe peritonitis where the cavity does not support expression of F4/80 that, given the dependence of F4/80 expression by LPM on GATA6 and retinoic acid^5,21,22^ **(Figure 1l)**, suggests severe peritoneal inflammation leads to a protracted but ultimately transient loss in retinoic acid availability.

### Inflammation leads to prolonged impairment of B1 cell accumulation in the peritoneal cavity

Although the failure to produce CXCL13 was a common feature of IMac-derived LPM **(Figure 4e)**, treatment with high-dose zymosan led to a striking reduction of CXCL13^+^ peritoneal macrophages **(Figure 4j)** arising from the comprehensive loss of the incumbent CXCL13-expressing resident cells. Given the non-redundant role of CXCL13 in maintenance of the peritoneal B1-cell pool^44^, we investigated whether peritoneal inflammation led to long-term disruption of B1 cells. Temporal analysis revealed that while the number of CD11b^+^ B1 cells gradually increased with age under homeostatic conditions **(Figure 4k)**, the degree of accumulation was slightly reduced following mild inflammation and completely abrogated following severe inflammation, yet neither led to absolute loss of B1 cells over baseline levels **(Figure 4l)**. Direct comparison of numbers of B1 cells in recipient mice from adoptive transfer experiments confirmed that severe inflammation led to substantially fewer CD11b^+^ peritoneal B1 cells within the cavity than following mild inflammation **(Supplementary Figure 5g)**. Furthermore, severe inflammation led to increased levels of serum IgM against phosphorylcholine, the predominant target of natural antibodies produced by peritoneal B1 cells^44,45^ and to the appearance of anti-phosphorylcholine IgG **(Figure 4m)**. Hence, sterile peritoneal inflammation leads to a state of altered homeostasis characterized by a failure to increase numbers of peritoneal CD11b^+^ B1 cells over time but which is associated with increased levels and class-switching of circulating natural antibody.

## Discussion

Transient peritoneal inflammation has lasting consequences for the incidence and severity of future disease^38,46^ but the mechanisms underlying this effect remain largely unresolved. Here, we demonstrate that inflammatory peritoneal macrophages recruited following sterile inflammation persist long-term but in an aberrant state of activation largely due to an inability to compete with incumbent macrophages for ‘niche’ signals and inflammation-driven alterations in the peritoneal environment. In so doing, we reveal the existence of multiple overlapping biochemical ‘niches’ that control programming of resident peritoneal macrophages and which are distinct from that controlling cell survival.

Like Liu et al^35^, our study suggests that the degree of replacement of LPM from the bone marrow following inflammation depends on the magnitude of initial macrophage disappearance. Furthermore, the increased survival of inflammatory macrophages following transfer into macrophage-depleted recipients provides definitive evidence that incumbent LPM impair survival of recruited cells. Hence, even without a defined physical niche, monocyte contribution to resident macrophages within fluidic environments appears subject to the same parameters of niche access and availability proposed by Guilliams and Scott^16^.

Likewise, our data support a model whereby competition for access to a ‘biochemical’ niche plays a critical role in determining the long-term transcriptional identity of inflammation-elicited macrophages. Specifically, inflammation-elicited macrophages that survived following mild inflammation exhibited striking long-term differences to incumbent resident macrophages including high MHCII and low GATA6 expression, yet more rapidly adopted a GATA-6^hi^ MHCII^lo^ resident-like phenotype and transcriptome following transfer into naive macrophage-depleted mice. The failure of inflammation-elicited macrophages to down-regulate MHCII expression following transfer into intact naïve mice confirms this feature arises from competition with incumbent resident macrophages for signals that regulate MHCII expression. Consistent with this, inflammation-elicited macrophages rapidly downregulated MHCII *in vitro* in response to RA-independent omental factors. Similarly, the GATA6^lo^ phenotype of inflammation-elicited macrophages following mild inflammation likely arises due to competition with enduring resident macrophages for retinoic acid, since more rapid acquisition of a GATA6^hi^ phenotype occurred following severe inflammation concurrent with the ablation of resident macrophages. Critically, inflammation-elicited macrophages gradually adopted features seemingly regulated by competition, suggesting that with time these cells receive sufficient cues to acquire a mature resident phenotype.

Several features of inflammation-elicited macrophages appeared regulated by changes in the cavity microenvironment post-inflammation. For example, severe inflammation led to sustained expression of MHCII by inflammation-elicited macrophages despite natural ablation of competing resident cells. Furthermore, the severely inflamed environment triggered MHCII expression by adoptively transferred resident LPM that would otherwise remain largely MHCII^−^ in a non-inflamed or mild inflammation setting. Hence, it seems likely severe inflammation triggers release of novel signals stimulatory for MHCII.

A small number of genes remained differentially expressed in inflammation-elicited macrophages following transfer into macrophage-depleted mice, supporting the notion that developmental origin influences macrophage identity^47^. These included features reprogrammed over time (eg Timd4, VSIG4, Cd209b), and permanent ‘legacy’ features of monocyte-derived cells (eg Sema4a, CD62L). The processes regulating these traits remains unclear^47^. However, the rapid acquisition of Tim4 expression by inflammation-elicited LPM following severe inflammation and the inhibition of this by transfer of Tim4^+^ resident macrophages provides proof-of-principle that seemingly origin-related time-dependent features of resident macrophages can be rapidly reprogrammed by appropriate niche signals.

While the panel of genes assessed here was relatively limited, our findings that gene expression is largely dictated by competition with resident cells or the post-inflammatory environment are likely to hold true on the transcriptome as a whole. For example, the overlap between environment-dependent genes and GATA-6 regulated genes^5,21,22^ suggests lower GATA6 expression by inflammation-elicited macrophages controls a significant proportion of their unique transcriptional profile. Similarly, the retarded expression of GATA6 by monocyte-derived LPM macrophages recruited during homeostasis also likely underlies a significant degree of the distinct transcriptional clustering of these cells. Critically, these data reveal that GATA6 expression does not act binarily but rather the level of expression has a critical role in determining LPM identity, as predicted for transcription factors with many competing target sites^48^.

One of our most intriguing findings was the difference in proliferative activity of incumbent resident and inflammation-elicited LPM, with only the latter exhibiting the capacity to overtly expand following transfer into macrophage-depleted mice. Notably, GATA6 directly regulates proliferation of LPM^22^, suggesting this disparity may relate to differences in GATA6 expression. However, treatment with exogenous CSF1 or IL-4 stimulates recently-recruited and incumbent resident macrophages to proliferate to equivalently high degrees^20,35,49^ and hence, the poor expansion of incumbent resident macrophages within the macrophage-deplete environment is not due to an intrinsic inability to proliferate.

Despite differentiating under inflamed conditions, persistent inflammation-elicited LPM bore striking similarities to monocyte-derived LPM present under non-inflamed conditions. As well as gene expression and proliferative activity, monocyte-derived LPM exhibited a largely comparable response to LPS irrespective of condition of differentiation, most notably characterized by increased production of IL-10 compared to embryonically-seeded LPM. Other than IL-1β, the profile of cytokine production by monocyte-derived LPM was the opposite reported for GATA6-deficient LPM^50^, suggesting other factors control their differential response to LPS. Hence, mild peritonitis does not lead to the existence of a unique subset of LPM but rather the expansion of a subset present in homeostasis. As we have previously shown that the abundance of Tim4^−^ monocyte-derived LPM gradually increases with age^20^, it would appear that mild inflammation accelerates a process normally associated with aging.

Our data also predict that inflammation-elicited LPM fail to express *Cxcl13* due to inflammation-induced loss of requisite niche signals. Notably, *Cxcl13* expression by LPM is sustained *in vitro* without addition of ‘niche’ factors^41^, potentially explaining why CXCL13 expression remains intact in incumbent LPM following mild inflammation. Hence, unlike the reversible programme of gene expression controlled by GATA6 that is lost in the absence of RA^41^, niche signals required to induce expression of CXCL13 in newly recruited macrophages may not be needed to maintain expression in resident cells.

CXCL13-deficient mice are profoundly deficient in peritoneal B1 cells^44^ yet CXCL13 is not required for retention of B1 cells in the cavity^51^. We found the extent of inflammation and consequently the ratio of CXCL13-producing resident macrophages to monocyte-derived CXCL13^−^ LPM in the cavity correlates with a failure to accumulate peritoneal B1 cell with age. As no other peritoneal lavage cells produce CXCL13^15,44^, our data suggest CXCL13 production by peritoneal macrophages is required for continued recruitment of B1 cells from the circulation. Notably, replacement of peritoneal LPM by monocytes and concurrent failure to expand peritoneal B1 cells also occurs following abdominal surgery^15^. Hence, long-term dysregulation of B1 cells is likely a general feature of peritoneal inflammation. Sterile peritoneal inflammation also led to elevated circulating natural antibody. Whereas splenic and bone marrow B1 cells spontaneously secrete high levels of natural IgM, those in the cavities do not^52^. Furthermore, levels of serum anti-PC IgM gradually drop with age^53^. Hence, we speculate that the failure to accumulate B1 cells in the peritoneal cavity may allow their re-entry into tissues permissive for antibody secretion such as fat associated lymphoid clusters^54^.

Inflammation-driven integration of functionally distinct monocyte-derived LPM is likely to occur in human peritoneal pathologies, as key features identified here are similar to published work on human peritoneal macrophages. Critically, human peritoneal macrophages are also considered Gata6-regulated, with 80% expressing detectable levels of this transcription factor ^55^. Stengel et al^56^ found rapid loss of CD206^+^LPM in response to spontaneous bacterial peritonitis consistent with a macrophage disappearance reaction which could allow for monocyte colonization. Indeed, two subsets of peritoneal macrophages exist in peritoneal ascites fluid from patients with decompensated cirrhosis, a more phagocytic subset expressing high levels of VSIG4 and Tim4 and a second less phagocytic subset exhibiting low levels of VSIG4, high levels of CCR2 and Sema4a, and responsiveness to retinoic acid^57^.

Our study highlights how varying degrees of inflammation alter the peritoneal macrophage compartment long-term and consequently reshape peritoneal homeostasis, and implicates competition for niche signals, time-of-residency and alterations in niche as principal determinants of these phenomena. These findings have broad importance for our understanding of plasticity within the mononuclear phagocyte compartment. Furthermore, understanding the consequences of inflammation in the serous cavities has major implications for pathologies in which serous cavity macrophages play key roles, including endometriosis^58^, adhesions^59^, and repair and scarring of visceral organs^19,60^ and the myriad of diseases influenced by natural antibody^61^.

## Materials and Methods

### Animals and reagents

C57BL/6 CD45.2^+^ and congenic CD45.1^+^CD45.2^+^ mice were bred and maintained in specific pathogen-free facilities at the University of Edinburgh, UK. In some experiments, C57BL/6JCrl mice were purchased from Charles River, UK. Mice were sex matched in all experiments and used at 6-10wks of age at the start of the experiment. Experiments were permitted under license by the UK Home Office and were approved by the University of Edinburgh Animal Welfare and Ethical Review Body. Details of reagents can be found in **Supplementary Table 5**.

### Sterile peritoneal inflammation

To elicit sterile peritoneal inflammation, mice were injected with 10 or 1000μg of zymosan A (Sigma-Aldrich) suspended in 200μl Dulbecco’s PBS (Invitrogen), dPBS or left naïve as indicated. In some experiments, mice were injected intraperitoneally with 250 μl of 700nm PKH26-PCL in suspended in Diluent B (Sigma) 24hrs prior to zymosan. In some experiments, mice were injected intraperitoneally with 0.0625mg Clodronate liposomes (Liposoma) suspended in 250μl dPBS (Gibco). To elicit LPS-induced inflammation mice were injected intraperitoneally with 5μg of LPS (O111:B4, Sigma-Aldrich) suspended in 200μl dPBS.

### Tissue-protected BM chimeric mice

Eight week-old female C57BL/6J CD45.1^+^CD45.2^+^ or CD45.2^+^ C57BL/6J mice were exposed to a single dose of 9.5 Gy γ-irradiation under anaesthetic, with all but the hind legs of the animals protected by a 2 inch lead shield. Animals were subsequently given 2-5×10^6^ BM cells from female congenic CD45.2^+^ C57BL/6J or C57BL/6J CD45.1^+^CD45.2^+^ animals respectively by i.v. injection and then left for 8wks, or in one experiment for 26wks due to the COVID-19-pandemic, before injection of zymosan.

### Cell isolation & Flow cytometry

Mice were sacrificed by exposure to rising levels of CO2. The peritoneal cavity was lavaged with a total of 9 ml wash solution (dPBS containing 2mM EDTA,1mM HEPES) or 9ml culture solution (RPMI containing 1mM HEPES) if cells were used for subsequent cell culture experiments. In some experiments, blood was then taken from the inferior vena cava and serum isolated using Microtainer SST tubes (BD). Serum was frozen at −80°C before analysis by ELISA. For chimeric mice, blood samples were also taken on the day of necropsy by cardiac puncture or by cutting the carotid artery. Equal cell numbers where incubated at room temperature for 10 minutes with zombie aqua viability dye (BioLegend), followed by 10-minute incubation on ice with blocking buffer containing 10% mouse serum with 0.25 μg/ml anti CD16/CD32 (BioLegend). Cells where incubated with indicated antibodies (**Supplementary Table 5**) on ice for 30 minutes. Cells where washed with FACS buffer (2mM EDTA/0.5%BSA in PBS) and if applicable stained with streptavidin conjugated or secondary antibody. For intracellular staining, cells where fixed/ permeabilized using the Foxp3 staining buffer (eBioscience) according to the manufacturers protocol. For intracellular cytokine staining 1E6 peritoneal cells were incubated for 4.5 hours at 37°C in 200μl sterile RPMI containing Brefeldin A and Monensin (both 1:1000) in cell repellent 96 well plates (Greiner Bio-One) after which cells were washed once and stained on ice for extracellular and intracellular as described with one additional Fc blocking step (10 minutes on ice) after fixation. Samples were acquired using FACS LSRFortessa (BD) and analysed using Flowjo (Version 10.4.1,Treestar). For analysis doublets (on the basis of Forward scatter area vs height) and dead cells (ZombieAqua positive) were excluded. For cell sorting, cells where stained using the same protocol scaled accordingly DAPI was used as viability dye to ensure real time viability detection. For adoptive transfer studies and cell culture studies cells were sorted using a FACSFusion of FACSAria sorting system with a 100μm sort nozzle. Cells sorted for RNA extraction were sorted using a 70μm nozzle.

### Adoptive transfer of macrophages

Cells were kept on ice at all time and all steps were carried out in a laminar flow hood using sterile reagents. Cells were collected and stained as described above and where sorted using flow cytometry into the indicated populations. Post sort cells where pelleted (300g, 5 min at 4°C), resuspended in 200μl of dPBS and counted by Casy Counter (Scharfe). For short-term and long-term studies 1×10^5^ and 1×10^5^ cells of the indicated populations were transferred respectively. For transfer of RMac^naive^ into high dose zymosan-treated recipients, 4×10^5^ cells were used. To study responsiveness to LPS in vivo, 2.5×10^5^ cells of each of the indicated populations was transferred. If required, purified populations from multiple donor mice were pooled. For each experiment, purified cells were suspended in 200μl of dPBS and injected intraperitoneally into recipients.

### Nanostring assay

For each of the indicated donor populations 5000 cells of interest where sorted into 2μl of RLT (QIAGEN). Immediately after cells where: centrifuged at maximum speed 15 seconds, vortexed for 10 seconds and again centrifuged at maximum speed 15 seconds. Cells where stored at −80°C until analysis using the nCounter Myeloid innate immunity panel (Nanostring) according to the manufacturer’s instructions. Data was analysed using the nSolver advanced analysis package. Differential gene expression was determined by pairwise comparison of IMAC and RMac^naive^ to RMac^naive^. Benjamini Hochberg adjusted p-value <0.05 were considered differentially expressed. Figures were generated using R packages GGplot2, Pheatmap, EnhancedVolcano and GOplot.

### Gene set enrichment analysis (GSEA)

Gene set of GATA6 regulated genes was obtained by analyzing GSE56711, GSE37448 and GSE47049 using the GEO2R. Genes were considered GATA6 regulated if they were differentially expressed (p<0.05) in 2 out of 3 published GATA6^KO^ datasets^5,21,22^. The gene list was split into GATA6^KO^ up and downregulated gene sets. GSEA was carried out using GSEA desktop 4.1 (Broad Institute). RNA levels of genes in RMac^Z10^ and IMac^Z10^ were analyzed using default settings and 10.000 geneset permutations.

### Omentum factors production and treatment

Omentum factors were generated by culturing the omentum from naïve mice in 1ml of macrophage serum free media (GIBCO) for 5 days as described^5^ after which medium was collected, centrifuged at 300g and the supernatant collected and diluted in 1:2 in media. Peritoneal cells were collected 11 days post low dose zymosan were collected as described under sterile conditions and 5×10^5^ plated and incubated for 2 hours at 37°C in cell culture medium (RPMI, 10%FCS, 1% L-Glutamine and 1% Pen/strep supplemented with 20ng/ml CSF1) after which medium was aspirated and cells were incubated in 250μl cell culture medium with 250μl Omentum factors or macrophage serum free media with ATRA (Sigma, 1μm) or without for 24 hours. Then, medium was removed and plate was incubated with 5mM EDTA in ice cold PBS for 10 minutes on ice to collect cells. Wells were repeatedly washed with ice cold 5mM EDTA PBS and wells were inspected using a microscope to confirm negligible adherent cells remained. Cells were quantified and prepared for flow cytometry as described.

### Cytokine production assay

Cells were kept on ice at all time and all steps were carried out using sterile reagents in a laminar flow hood using the sort protocol described. For each population of interest 1×10^3^ cells per condition were sorted into 75μl sort medium (Folic acid deficient RPMI containing 20% FCS (Low endotoxin), 2% L-Glutamine and 2% Pen/strep). Cells were centrifuged at 100g at 4 degrees for 5 minutes. The total mixture was then transferred into a 96 well plate incubated at 37°C for 2 hours. Media was gently aspirated and cells were resuspended in 75μl cell culture medium (Folic acid deficient RPMI supplemented with 1μg/ml Folic Acid (Sigma-Aldrich), 10% FCS (Low endotoxin), 1% L-glutamine and 1% Pen/Strep). Where indicated cells received a final concentration of LPS of 1ng/ml (O11:B4, Sigma-Aldrich) in cell culture medium or equivalent amount of dPBS. Cells were incubated for 14 hours and supernatant was collected and analysed for cytokine release using the Legendplex Mouse Anti-Virus or Mouse Inflammation panel according to the manufacturers protocol. Data was acquired using an Attune flow cytometer and analysed using the Legendplex analysis software.

### Phrodo phagocytosis assay

Cells were collected and 2×10^6^ cells/sample were stained as described. Each sample was washed twice with ice cold RPMI and was split into two tubes each and left on ice for 10 minutes. Then to each tube 10μl of Phrodo E.Coli particles was added and for each sample one tube was incubated at 37°C and one at 4°C for 1hr. All samples were placed on ice and washed once using 300μl Buffer C and were then resuspended in 300μl Buffer C. Cells were analysed directly after finishing the protocol. Data is presented as normalized Phrodo mean fluorescence intensity (MFI 37°C-MFI 4°C).

#### Enzyme-Linked Immunosorbant assay’s

96-well flat-bottom high-binding polystyrene plates (Corning) were coated with 50μl of 2μg/ml phosphorylcholine conjugated to BSA (PC-BSA; 2B Scientific) diluted in PBS at 4°C overnight. Plates were then blocked with 100μl of blocking buffer (1% Casein in PBS; VWR) for 1.5hr at room temperature, before serum samples were added at 1:100 dilution in 50μl blocking buffer and incubated for 2hr at room temperature. Wells without antigen were used as blank controls for each sample to measure non-specific antibody binding. Plates were then incubated with 1:5000 HRP-conjugated anti-mouse IgG (Abcam) or 1:2000 anti-mouse IgM (Southern Biotech) in blocking buffer for 1h at room temperature before addition of TMB (Seracare). After 10 minutes the reaction was stopped with 0.16M sulphuric acid solution and the OD450 value measured. Values for blank controls were then subtracted for each sample to quantify antigen-specific antibody levels. Plates were washed twice with 0.1% Tween20 (Sigma-Aldrich) in PBS between all steps except before addition of TMB, when they were washed 5 times.

#### Statistics

Statistics were performed using Prism 7 (GraphPad Software). The statistical test used in each experiment is detailed in the relevant figure legend.

#### Accession codes

Nanostring gene expression data that support the findings of this study have been deposited in -------

#### Data availability

Data that support the findings of this study are available from the corresponding authors upon reasonable request.

## Acknowledgements

Flow cytometry data were generated with support from the QMRI Flow Cytometry and Cell Sorting Facility, University of Edinburgh. Nanostring was performed by the Host and Tumour Profiling Unit Services Facility, MRC Institute for Genetics and Molecular Medicine, University of Edinburgh. We thank Dr Mohini Gray for kind provision of PC-BSA and Prof Judith Allen for comments on the manuscript.

This work was funded by a Wellcome Trust PhD studentship to P.A.L (203909/Z/16/A) with additional support from the Medical Research Council UK (MR/L008076/1 to S.J.J).

## Author Contributions

P.A.L. designed and performed most experiments, analysed and interpreted the data, and wrote the manuscript. L.B.G. performed and analysed antibody ELISA’s. H.W and G.P-W. performed experiments. C.C.B contributed to design of experiments and interpretation of data. S.J.F. provided critical feedback on the study design. S.J.J. conceived, designed and performed experiments, analysed and interpreted the data, wrote the manuscript, and supervised the project.

---

**Supplementary Figure 1.**
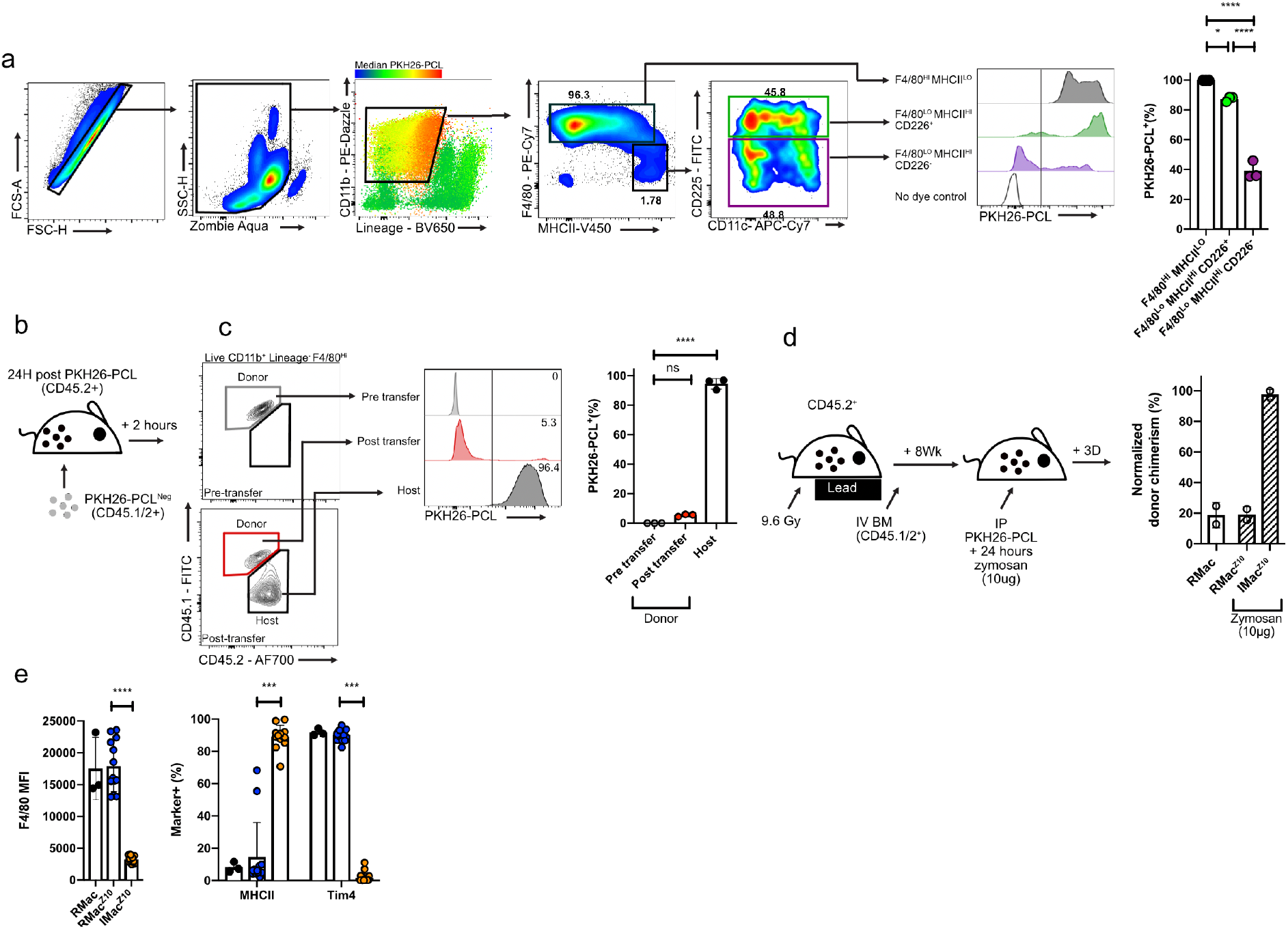
A toolbox to investigate inflammatory macrophage fate. **(a)** Dye labelling efficiency of F4/80^HI^ MHCII^LO^ resident macrophages (grey) and F4/80^LO^MHCII^HI^ CD226^+^ small peritoneal macrophages (green) or CD226^−^ DCs or immature macrophages (purple) 24hrs after intraperitoneal administration of PKH26-PCL (n=3). **p<0.01 by determined one-way ANOVA with Tukey’s multiple comparisons test. **(b)** Experimental scheme for the adoptive transfer of unlabelled CD45.1/2^+^ peritoneal exudate cells (PEC) into the peritoneal cavity of CD45.2^+^ mice injected with PKH26-PCL intraperitoneally 24hrs prior. **(c)** Representative PKH26-PCL labelling and quantification of donor F4/80^hi^ macrophages prior to transfer (top) and 2hrs post transfer (red; n=3) compared to recipient F4/80^Hi^ macrophages (black). ****p<0.0001 determined by one-way ANOVA with Tukey’s multiple comparisons test. **(d)** Non-host chimerism of F4/80^HI^ PKH26-PCL^HI^ RMac in the naïve peritoneal cavity (white bar) and F4/80^HI^ PKH26-PCL^HI^ RMac^Z10^ and PKH26-PCL^LO^ F4/80^INT^ IMac^Z10^ 3d post 10μg zymosan (hatched bars). Dye injection given 8wks after irradiation and zymosan injection given 24hrs thereafter. **(e)** Expression of F4/80, MHCII and Tim4 by RMac(black), RMac^Z10^(blue) and IMac^Z10^ 3 days post 10μg zymosan. ***p<0.001 ****p<0.0001 determined one-way ANOVA with Dunnet’s multiple comparisons test for each marker individually, followed by Bonferroni adjustment. For all experiments, data are presented as mean ± standard deviation with each symbol representing an individual animal. All data were pooled from at least 2 independent experiments, except for **(a,c,f)** which are from single experiments.

**Supplementary Figure 2.**
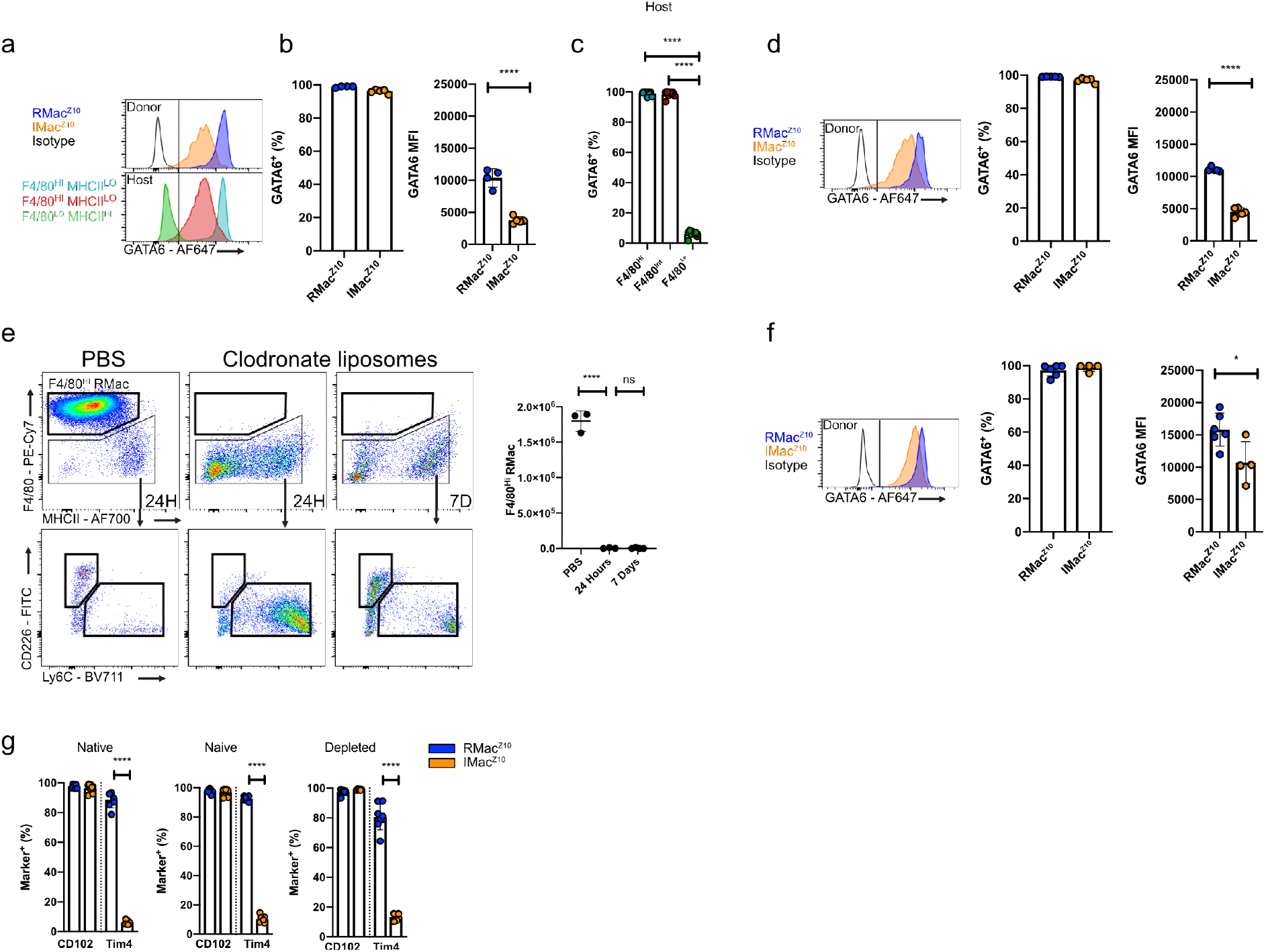
Competition mediates inflammatory macrophage survival and phenotype during early resolution. **(a)** Representative expression of GATA6 by indicated donor populations 8d post transfer into inflamed recipients (top) and by host macrophage subsets identified using F4/80 and MHCII (bottom). **(b)** Proportion of donor RMac^Z10^(n=4), IMac^Z10^ (n=5) that were GATA6^+^ and MFI of GATA6 expression 8d after transfer into inflamed recipients. ****p<0.0001 determined by student’s t test. **(c)** Proportion of host macrophage subsets that are GATA6^+^ 11d post zymosan (8d post cell transfer; n=9) ****p<0.0001 determined by one-way ANOVA with Tukey’s multiple comparisons test. **(d)** Proportion of donor RMac^Z10^ (n=5) and IMac^Z10^ (n=5) that were GATA6^+^ and MFI of GATA6 expression 8d after transfer into naive recipients. ****p<0.0001 determined by student’s t test. **(e)** Representative dot-plots gated on CD11b^+^ cells and absolute number of F4/80^Hi^ resident macrophages (black) 24hrs after intraperitoneal injection of PBS (n=3) or clodronate liposomes (n=3) and 7d post clodronate liposome injection (n=4). ****p<0.0001 determined by one-way ANOVA and Dunnet’s multiple comparisons test. **(f)** Proportion of donor RMac^Z10^ (n=6) and IMac^Z10^ (n=5) that were GATA6^+^ and MFI of GATA6 expression 8d after transfer into clodronate depleted recipients. *p<0.05 determined by student’s t test. **(g)** Proportion of donor RMac^Z10^ and IMac^Z10^that are CD102^+^ and Tim4^+^ 8d after transfer into mirroring native (n= 7, 8), naïve (n=7) or clodronate depleted recipients (n= 8,6). ****p<0.0001 determined by one-way ANOVA and Sidak’s multiple comparisons test. For all experiments, data are presented as mean ± standard deviation with each symbol representing an individual animal. All data were pooled from at least 2 independent experiments, except for **(e)** is from a single experiment.

**Supplementary Figure 3.**
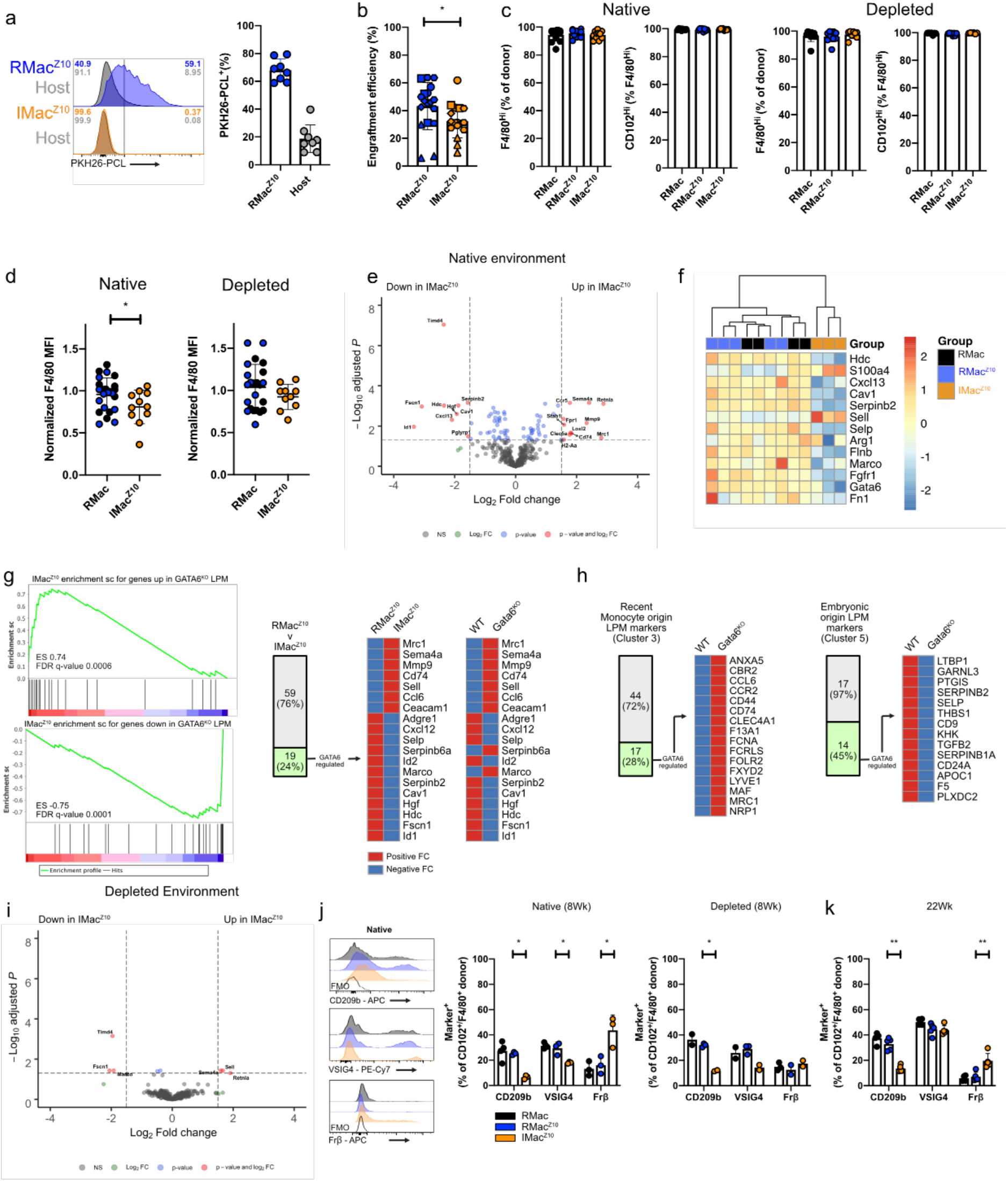
Colonizing inflammatory macrophages are long lived but retain cell intrinsic and environment dependent transcriptional and phenotypic differences. **(a)** Proportion of donor CD45.2^+^ F4/80^Hi^ RMac^Z10^and host CD45.1/2^+^ F4/80^Hi^ macrophages that are PKH26-PCL^+^ 8wks post transfer (n=8). **(b)** pooled engraftment efficiency of RMac^Z10^ and IMac^Z10^8wks post transfer into native recipients. Includes data presented in Figure 2b and Figure 4b. Each symbol refers to an experimental run. *p<0.05 determined by student’s t test. **(c)** Fraction of donor RMac, RMac^Z10^ and IMac^Z10^ that are F4/80^Hi^ and the proportion of which are CD102^Hi^ after transfer into native (left; n= 12,11,11) and depleted recipients (right; n = 10,8,8). **(d)** Normalized F4/80 MFI on donor RMac(black), RMac^Z10^ (blue) and IMac^Z10^ (orange) after transfer into native (left; n=12,11,11) or depleted (right; n=10,10,8) recipients. F4/80 MFI is normalized to mean F4/80 MFI of RMac. *p<0.05 determined by student’s t test **(e)** Volcano plot of gene expression of IMac^Z10^ relative to RMac^Z10^ 8wks post transfer into native recipients. **(f)** Heatmap highlighting the subset of peritoneal macrophage identity genes included in the Nanostring panel and their expression by donor RMac, RMac^Z10^ and IMac^Z10^ 8wks post transfer into native recipients. **(g)** GSEA of mrNA in RMac^Z10^and IMac^Z10^ against genes up and downregulated genes in GATA6^KO^ LPM (left). In the middle, proportion of differentially expressed genes between IMac^Z10^ and RMac^Z10^ that are regulated by GATA6 (left) and their transcriptional directionality. On the right, transcriptional directionality of the same genes in GATA6^KO^ LPM relative to WT. **(h)** Proportion of highly expressed single cell cluster genes that are regulated by GATA6 in cluster 3 and 5 as described by Bain et al^15^ and the transcriptional directionality of these genes in GATA6^KO^ LPM relative to WT. **(i)** Volcano plot of gene expression of IMac^Z10^ relative to RMac^Z10^ 8wks post transfer into depleted recipients. **(j)** Expression of markers of interest by CD102^+^ or F4/80^+^ donor RMac (black) RMac^Z10^ (blue), IMac^Z10^ (orange) 8wks post transfer into native (n=4,3,3) and depleted recipients (n=2,3,2). Data obtained from a single experiment. *p<0.05 determined by one way Anova and Dunnet’s multiple comparisons test for each marker individually, followed by Bonferroni adjustment. **(k)** Expression of markers of interest by CD102^+^ or F4/80^+^ donor RMac(black; n=4) RMac^Z10^ (blue; n=5), IMac^Z10^ (orange; n=5) 22wks post transfer into native recipients. **p<0.01 determined by one way Anova and Dunnet’s multiple comparisons test for each marker individually, followed by Bonferroni adjustment. For all experiments, data are presented as mean ± standard deviation with each symbol representing an individual animal. All data were pooled from at least 2 independent experiments, except for **(i)** which is from a single experiments.

**Supplementary Figure 4.**
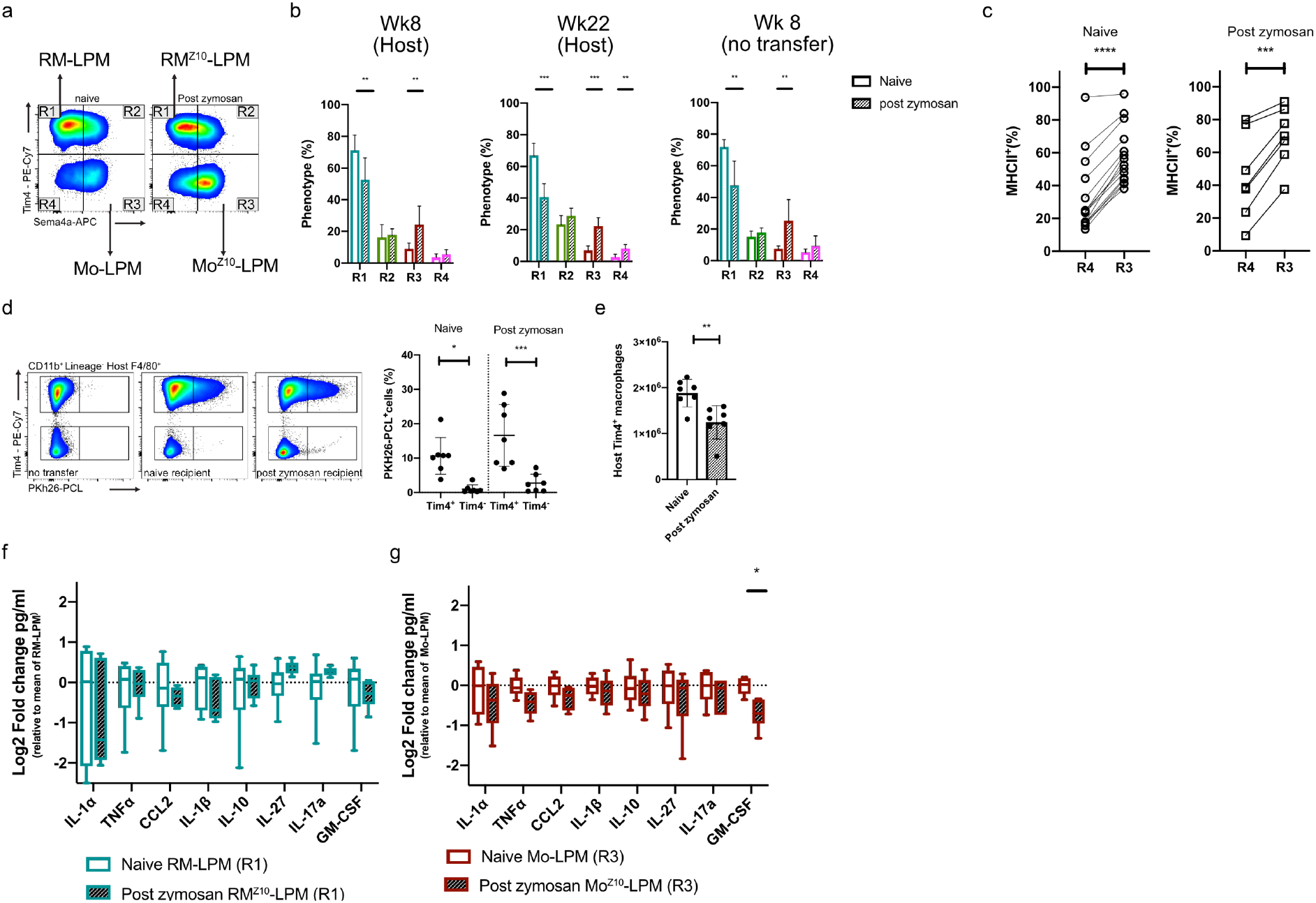
Monocyte derived LPM are functionally distinct from embryonically seeded LPM. **(a)** Representative expression of Tim4 and Sema4a by LPM from naïve mice or 8wks post zymosan injection. **(b)** Proportion of host LPM with Sema4a^LO^Tim4^+^(R1), Sema4a^HI^Tim4^+^(R2), Sema4a^Hi^Tim4^−^ (R3) or Sema4a^Lo^Tim4^−^ (R4) phenotype 8wks post zymosan (6 naïve v 12 zymosan treated) and 22wks post zymosan 4 naïve v 10 zymosan treated) compared to naïve control. On the right, proportion of LPM with Sema4a^LO^Tim4^+^(R1), Sema4a^HI^Tim4^+^(R2), Sema4a^Hi^Tim4^−^ (R3) or Sema4a^Lo^Tim4^−^ (R4) phenotype 8wks post zymosan (15 naïve v 8 zymosan treated). ***p<0.001**p<0.01 determined by repeated student’s t test with Holm-Sidak correction. **(c)** Proportion of Sema4a^Lo^Tim4^−^ (R4) and Sema4a^Hi^Tim4^−^ (R3) that are MHCII^+^ in naïve (n=15) and zymosan treated (n=7) mice. ***p<0.001****p<0.0001 determined by paired student’s t test. **(d)** Fraction of naïve or post zymosan host F4/80^+^Tim4^+/-^ macrophages that are PKH26-PCL labelled 8d after receiving PKH26-PCL labelled RMac(n=7). *p<0.05***p<0.0001 determined by one way ANOVA with Sidak multiple comparison test. **(e)** Absolute number of host naïve or post zymosan host Tim4^+^ macrophages 8d after receiving PKH26-PCL labelled RMac (n=7). **p<0.01 determined by student’s t test. **(f)** Secreted cytokine/chemokine profile collected from cultures of RM-LPM or RM^Z10^-LPM (n=6), sourced from naïve or 8wks post zymosan mice, 14hrs after culture with LPS (1ng/ml). Results are shown as log2 fold change in mean pg/ml over the mean RM-LPM using a box-and-whiskers plot. **(g)** Secreted cytokine/chemokine profile collected from cultures of Mo-LPM or Mo^Z10^-LPM (n=6), sourced from naïve or 8wks post zymosan mice, 14hrs after culture with LPS (1ng/ml). Results are shown as log2 fold change in mean pg/ml over the mean Mo-LPM using a box-and-whiskers plot. *p<0.05determined by repeated student’s t test with Holm-Sidak correction. For all experiments, data are presented as mean ± standard deviation with each symbol representing an individual animal. All data were pooled from at least 2 independent experiments.

**Supplementary Figure 5.**
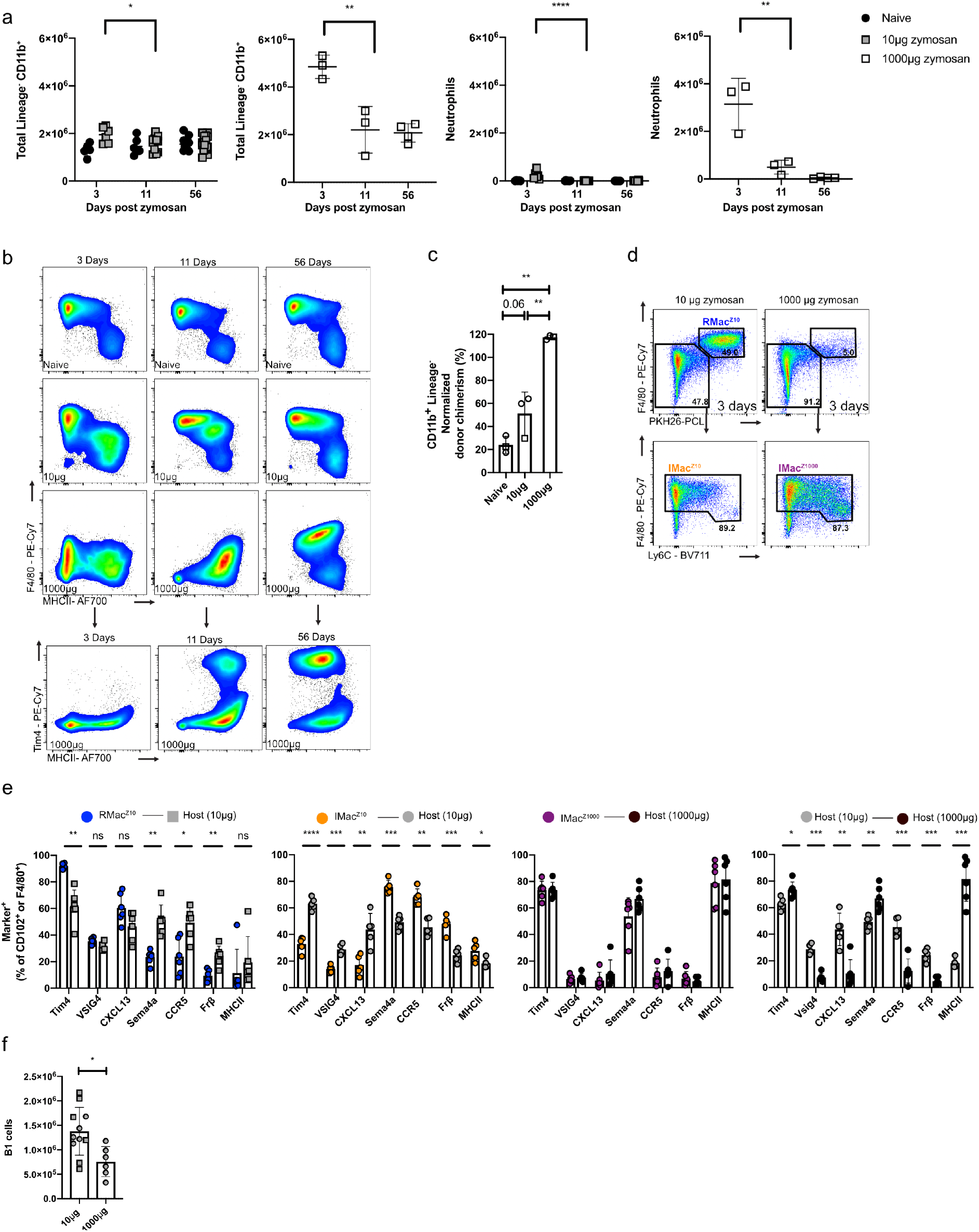
Ontogeny does not dictate monocyte phenotype after severe peritonitis in females and leads to impaired B1 cell expansion. **(a)** Absolute number of CD11b^+^ myeloid cells and neutrophils at indicated timepoints in naïve (black circle; n= 5,6,10), post low dose zymosan (grey square; n=6,11,20) and post high dose zymosan (white square; n=3,3,4) *p<0.05, ***p<0.001, ****p<0.0001 for naïve and low dose determined by two-way ANOVA and post hoc Tukey test. For high dose determined by one-way ANOVA and post hoc Tukey test **(b)** Representative expression of F4/80, MHCII and Tim4 by at indicated timepoints in naïve, 10 or 1000μg zymosan treated mice. **(c)** Non host chimerism of CD11B^+^ myeloid cells 17d after indicated zymosan dose in tissue-protected BM chimeric mice. Zymosan treatment 8 (circle) or 26 (square) wks after irradiation. **p<0.01 determined by one-way ANOVA and Tukey’s multiple comparisons test **(d)** Representative expression of F4/80, Ly6C and PKH26-PCL labelling and identification of F4/80^HI^ PKH26-PCL^HI^ resident macrophages (RMac^Z10^) and PKh26-PCL^LO^ F4/80^INT^ inflammatory macrophages3d after 10μg (IMac^Z10^) or 1000μg (IMac^Z1000^) zymosan. **(e)** Expression of markers of interest by CD102^+^/F4/80^+^ donor RMac^Z10^(n=6), IMac^Z10^ (n=5) or IMac^Z1000^ (n=6) and their respective CD102^+^/F4/80^+^ host macrophages. *p<0.05 **p<0.01 ***p<0.001 ****p<0.001 determined by repeated student’s t test with Holm-Sidak correction **(f)** Absolute number of CD11b^+^ B1 cells 8wks after 10μg of zymosan (n=11) or 1000μg zymosan (n=6) with transfer or IMac or IMac^1000^ at D3 respectively. *p<0.05 determined by student’s t test. For all experiments, data are presented as mean ± standard deviation with each symbol representing an individual animal. All data were pooled from at least 2 independent experiments except **(a)** where high dose datapoints where from a single experiment.

## References

1 Davies, L. C., Jenkins, S. J., Allen, J. E. & Taylor, P. R. Tissue-resident macrophages. Nature immunology 14, 986–995, doi:10.1038/ni.2705 (2013).

2 Gautier, E. L. et al. Gene-expression profiles and transcriptional regulatory pathways that underlie the identity and diversity of mouse tissue macrophages. Nature immunology 13, 1118–1128, doi:10.1038/ni.2419 (2012).

3 Gosselin, D. et al. Environment drives selection and function of enhancers controlling tissue-specific macrophage identities. Cell 159, 1327–1340, doi:10.1016/j.cell.2014.11.023 (2014).

4 Lavin, Y. et al. Tissue-resident macrophage enhancer landscapes are shaped by the local microenvironment. Cell 159, 1312–1326, doi:10.1016/j.cell.2014.11.018 (2014).

5 Okabe, Y. & Medzhitov, R. Tissue-specific signals control reversible program of localization and functional polarization of macrophages. Cell 157, 832–844, doi:10.1016/j.cell.2014.04.016 (2014).

6 Ginhoux, F. & Guilliams, M. Tissue-Resident Macrophage Ontogeny and Homeostasis. Immunity 44, 439–449, doi:10.1016/j.immuni.2016.02.024 (2016).

7 Hashimoto, D. et al. Tissue-resident macrophages self-maintain locally throughout adult life with minimal contribution from circulating monocytes. Immunity 38, 792–804, doi:10.1016/j.immuni.2013.04.004 (2013).

8 De Schepper, S. et al. Self-Maintaining Gut Macrophages Are Essential for Intestinal Homeostasis. Cell 175, 400–415 e413, doi:10.1016/j.cell.2018.07.048 (2018).

9 Sierro, F. et al. A Liver Capsular Network of Monocyte-Derived Macrophages Restricts Hepatic Dissemination of Intraperitoneal Bacteria by Neutrophil Recruitment. Immunity 47, 374–388 e376, doi:10.1016/j.immuni.2017.07.018 (2017).

10 Beattie, L. et al. Bone marrow-derived and resident liver macrophages display unique transcriptomic signatures but similar biological functions. J Hepatol 65, 758–768, doi:10.1016/j.jhep.2016.05.037 (2016).

11 David, B. A. et al. Combination of Mass Cytometry and Imaging Analysis Reveals Origin, Location, and Functional Repopulation of Liver Myeloid Cells in Mice. Gastroenterology 151, 1176–1191, doi:10.1053/j.gastro.2016.08.024 (2016).

12 Gibbings, S. L. et al. Transcriptome analysis highlights the conserved difference between embryonic and postnatal-derived alveolar macrophages. Blood 126, 1357–1366, doi:10.1182/blood-2015-01-624809 (2015).

13 Scott, C. L. et al. Bone marrow-derived monocytes give rise to self-renewing and fully differentiated Kupffer cells. Nat Commun 7, 10321, doi:10.1038/ncomms10321 (2016).

14 van de Laar, L. et al. Yolk Sac Macrophages, Fetal Liver, and Adult Monocytes Can Colonize an Empty Niche and Develop into Functional Tissue-Resident Macrophages. Immunity 44, 755–768, doi:10.1016/j.immuni.2016.02.017 (2016).

15 Bain, C. C. et al. Rate of replenishment and microenvironment contribute to the sexually dimorphic phenotype and function of peritoneal macrophages. Sci Immunol 5, doi:10.1126/sciimmunol.abc4466 (2020).

16 Guilliams, M. & Scott, C. L. Does niche competition determine the origin of tissue-resident macrophages? Nature reviews. Immunology 17, 451–460, doi:10.1038/nri.2017.42 (2017).

17 Zeng, Z. et al. Sex-hormone-driven innate antibodies protect females and infants against EPEC infection. Nature immunology 19, 1100–1111, doi:10.1038/s41590-018-0211-2 (2018).

18 Bain, C. C. & Jenkins, S. J. The biology of serous cavity macrophages. Cellular immunology, doi:10.1016/j.cellimm.2018.01.003 (2018).

19 Wang, J. & Kubes, P. A Reservoir of Mature Cavity Macrophages that Can Rapidly Invade Visceral Organs to Affect Tissue Repair. Cell 165, 668–678, doi:10.1016/j.cell.2016.03.009 (2016).

20 Bain, C. C. et al. Long-lived self-renewing bone marrow-derived macrophages displace embryo-derived cells to inhabit adult serous cavities. Nat Commun 7, ncomms11852, doi:10.1038/ncomms11852 (2016).

21 Gautier, E. L. et al. Gata6 regulates aspartoacylase expression in resident peritoneal macrophages and controls their survival. The Journal of experimental medicine 211, 1525–1531, doi:10.1084/jem.20140570 (2014).

22 Rosas, M. et al. The transcription factor Gata6 links tissue macrophage phenotype and proliferative renewal. Science 344, 645–648, doi:10.1126/science.1251414 (2014).

23 Cain, D. W. et al. Identification of a tissue-specific, C/EBPbeta-dependent pathway of differentiation for murine peritoneal macrophages. J Immunol 191, 4665–4675, doi:10.4049/jimmunol.1300581 (2013).

24 Kim, K. W. et al. MHC II+ resident peritoneal and pleural macrophages rely on IRF4 for development from circulating monocytes. The Journal of experimental medicine 213, 1951–1959, doi:10.1084/jem.20160486 (2016).

25 Casanova-Acebes, M. et al. RXRs control serous macrophage neonatal expansion and identity and contribute to ovarian cancer progression. Nat Commun 11, 1655, doi:10.1038/s41467-020-15371-0 (2020).

26 Zhou, X. et al. Circuit Design Features of a Stable Two-Cell System. Cell 172, 744–757 e717, doi:10.1016/j.cell.2018.01.015 (2018).

27 Bonnardel, J. et al. Stellate Cells, Hepatocytes, and Endothelial Cells Imprint the Kupffer Cell Identity on Monocytes Colonizing the Liver Macrophage Niche. Immunity 51, 638–654 e639, doi:10.1016/j.immuni.2019.08.017 (2019).

28 Chakarov, S. et al. Two distinct interstitial macrophage populations coexist across tissues in specific subtissular niches. Science 363, doi:10.1126/science.aau0964 (2019).

29 Mondor, I. et al. Lymphatic Endothelial Cells Are Essential Components of the Subcapsular Sinus Macrophage Niche. Immunity 50, 1453–1466 e1454, doi:10.1016/j.immuni.2019.04.002 (2019).

30 Zhang, N. et al. Expression of factor V by resident macrophages boosts host defense in the peritoneal cavity. The Journal of experimental medicine 216, 1291–1300, doi:10.1084/jem.20182024 (2019).

31 Gautier, E. L., Ivanov, S., Lesnik, P. & Randolph, G. J. Local apoptosis mediates clearance of macrophages from resolving inflammation in mice. Blood 122, 2714–2722, doi:10.1182/blood-2013-01-478206 (2013).

32 Zigmond, E. et al. Ly6C hi monocytes in the inflamed colon give rise to proinflammatory effector cells and migratory antigen-presenting cells. Immunity 37, 1076–1090, doi:10.1016/j.immuni.2012.08.026 (2012).

33 Aegerter, H. et al. Influenza-induced monocyte-derived alveolar macrophages confer prolonged antibacterial protection. Nature immunology 21, 145–157, doi:10.1038/s41590-019-0568-x (2020).

34 Misharin, A. V. et al. Monocyte-derived alveolar macrophages drive lung fibrosis and persist in the lung over the life span. The Journal of experimental medicine 214, 2387–2404, doi:10.1084/jem.20162152 (2017).

35 Liu, Z. et al. Fate Mapping via Ms4a3-Expression History Traces Monocyte-Derived Cells. Cell 178, 1509–1525 e1519, doi:10.1016/j.cell.2019.08.009 (2019).

36 Davies, L. C. et al. Distinct bone marrow-derived and tissue-resident macrophage lineages proliferate at key stages during inflammation. Nat Commun 4, 1886, doi:10.1038/ncomms2877 (2013).

37 Davies, L. C. et al. A quantifiable proliferative burst of tissue macrophages restores homeostatic macrophage populations after acute inflammation. European journal of immunology 41, 2155–2164, doi:10.1002/eji.201141817 (2011).

38 Newson, J. et al. Resolution of acute inflammation bridges the gap between innate and adaptive immunity. Blood 124, 1748–1764, doi:10.1182/blood-2014-03-562710 (2014).

39 Yona, S. et al. Fate Mapping Reveals Origins and Dynamics of Monocytes and Tissue Macrophages under Homeostasis. Immunity 38, 79–91, doi:10.1016/j.immuni.2012.12.001 (2013).

40 Liao, C. T. et al. IL-10 differentially controls the infiltration of inflammatory macrophages and antigen-presenting cells during inflammation. European journal of immunology 46, 2222–2232, doi:10.1002/eji.201646528 (2016).

41 Buechler, M. B. et al. A Stromal Niche Defined by Expression of the Transcription Factor WT1 Mediates Programming and Homeostasis of Cavity-Resident Macrophages. Immunity 51, 119–130 e115, doi:10.1016/j.immuni.2019.05.010 (2019).

42 Cash, J. L., White, G. E. & Greaves, D. R. Chapter 17. Zymosan-induced peritonitis as a simple experimental system for the study of inflammation. Methods Enzymol 461, 379–396, doi:10.1016/S0076-6879(09)05417-2 (2009).

43 Gundra, U. M. et al. Vitamin A mediates conversion of monocyte-derived macrophages into tissue-resident macrophages during alternative activation. Nature immunology 18, 642–653, doi:10.1038/ni.3734 (2017).

44 Ansel, K. M., Harris, R. B. & Cyster, J. G. CXCL13 is required for B1 cell homing, natural antibody production, and body cavity immunity. Immunity 16, 67–76 (2002).

45 Masmoudi, H., Mota-Santos, T., Huetz, F., Coutinho, A. & Cazenave, P. A. All T15 Id-positive antibodies (but not the majority of VHT15+ antibodies) are produced by peritoneal CD5+ B lymphocytes. Int Immunol 2, 515–520, doi:10.1093/intimm/2.6.515 (1990).

46 Newson, J. et al. Inflammatory Resolution Triggers a Prolonged Phase of Immune Suppression through COX-1/mPGES-1-Derived Prostaglandin E2. Cell Rep 20, 3162–3175, doi:10.1016/j.celrep.2017.08.098 (2017).

47 Bleriot, C., Chakarov, S. & Ginhoux, F. Determinants of Resident Tissue Macrophage Identity and Function. Immunity 52, 957–970, doi:10.1016/j.immuni.2020.05.014 (2020).

48 Brewster, R. C. et al. The transcription factor titration effect dictates level of gene expression. Cell 156, 1312–1323, doi:10.1016/j.cell.2014.02.022 (2014).

49 Jenkins, S. J. et al. Local macrophage proliferation, rather than recruitment from the blood, is a signature of TH2 inflammation. Science 332, 1284–1288, doi:10.1126/science.1204351 (2011).

50 Ipseiz, N. et al. Tissue-resident macrophages actively suppress IL-1beta release via a reactive prostanoid/IL-10 pathway. The EMBO journal 39, e103454, doi:10.15252/embj.2019103454 (2020).

51 Ha, S. A. et al. Regulation of B1 cell migration by signals through Toll-like receptors. The Journal of experimental medicine 203, 2541–2550, doi:10.1084/jem.20061041 (2006).

52 Choi, Y. S., Dieter, J. A., Rothaeusler, K., Luo, Z. & Baumgarth, N. B-1 cells in the bone marrow are a significant source of natural IgM. European journal of immunology 42, 120–129, doi:10.1002/eji.201141890 (2012).

53 Holodick, N. E., Vizconde, T., Hopkins, T. J. & Rothstein, T. L. Age-Related Decline in Natural IgM Function: Diversification and Selection of the B-1a Cell Pool with Age. J Immunol 196, 4348–4357, doi:10.4049/jimmunol.1600073 (2016).

54 Jackson-Jones, L. H. et al. Fat-associated lymphoid clusters control local IgM secretion during pleural infection and lung inflammation. Nat Commun 7, 12651, doi:10.1038/ncomms12651 (2016).

55 Ruiz-Alcaraz, A. J. et al. Characterization of human peritoneal monocyte/macrophage subsets in homeostasis: Phenotype, GATA6, phagocytic/oxidative activities and cytokines expression. Sci Rep 8, 12794, doi:10.1038/s41598-018-30787-x (2018).

56 Stengel, S. et al. Peritoneal Level of CD206 Associates With Mortality and an Inflammatory Macrophage Phenotype in Patients With Decompensated Cirrhosis and Spontaneous Bacterial Peritonitis. Gastroenterology 158, 1745–1761, doi:10.1053/j.gastro.2020.01.029 (2020).

57 Irvine, K. M. et al. CRIg-expressing peritoneal macrophages are associated with disease severity in patients with cirrhosis and ascites. JCI Insight 1, e86914, doi:10.1172/jci.insight.86914 (2016).

58 Hogg, C. D., P.; Rosser, M.; Mack, M.; Soong, D.; Pollard, J.W.; Jenkins, S.J.; Horne, A.W.; Greaves, E. Macrophages inhibit and enhance endometriosis depending on their origin. bioRxiv https://doi.org/10.1101/2020.04.30.070003 (2020).

59 Honjo, K. et al. Plasminogen activator inhibitor-1 regulates macrophage-dependent postoperative adhesion by enhancing EGF-HER1 signaling in mice. FASEB journal : official publication of the Federation of American Societies for Experimental Biology 31, 2625–2637, doi:10.1096/fj.201600871RR (2017).

60 Deniset, J. F. et al. Gata6(+) Pericardial Cavity Macrophages Relocate to the Injured Heart and Prevent Cardiac Fibrosis. Immunity 51, 131–140 e135, doi:10.1016/j.immuni.2019.06.010 (2019).

61 Hernandez, A. M. & Holodick, N. E. Editorial: Natural Antibodies in Health and Disease. Front Immunol 8, 1795, doi:10.3389/fimmu.2017.01795 (2017).

