## Supplementary Table 5 for "Recruited macrophages that colonise the post-inflammatory peritoneal niche convert into functionally divergent resident cells"

**Reagents used**

| **Reagent** |  | **Source** |  | **Catalogue #** |
| --- | --- | --- | --- | --- |
| BSA |  | Sigma-Aldrich |  | A7906-100g |
| Casein |  | VWR |  | 22544.292 |
| Clodronate Liposomes |  | Liposoma |  | n/a |
| dPBS |  | Gibco/ThermoFisher |  | 14190-094 |
| EDTA 0.5M |  | Invitrogen |  | 15574020 |
| FCS |  | Gibco/ThermoFisher |  | 10500-064 |
| Folic Acid |  | Sigma-Aldrich |  | F8758-5G |
| FSC (LE confirmed in house) |  | GE-Healthcare |  |  |
| HEPES 1M |  | Fisher Scientific |  | 10041703 |
| Intracellular Fixation and Permeabilization set |  | ThermoFisher |  | 889-8824-00 |
| L-Glutamine 200mM |  | Gibco/ThermoFisher |  | 25030024 |
| LEGENDplex mouse anti-virus |  | Biolegend |  | 740622 |
| LEGENDplex mouse inflammation |  | Biolegend |  | 740446 |
| LPS (O111:B4 |  | Sigma-Aldrich |  | L2630-10MG |
| LPS (O127:B8) |  | Sigma-Aldrich |  | L3129-10MG |
| Macrophage SFM |  | Gibco/ThermoFisher |  | 12065074 |
| PC-BSA |  | 2B Scientific |  | PC-1011-10 |
| Pen/Strep |  | Gibco/ThermoFisher |  | 15140122 |
| Phrodo E.coli phagocytosis kit |  | Invitrogen/ThermoFisher |  | A10025 |
| PKH26-PCL Cell Linker kit |  | Sigma-Aldrich |  | PKH26PCL-1KT |
| Recombinant murine CSF1 |  | Peprotech |  | 315-02 |
| Retinoic Acid |  | Sigma-Aldrich |  | R2625-50MG |
| RPMI 1640 |  | Gibco/ThermoFisher |  | 21870076 |
| RPMI 1640 no folic acid |  | Gibco/ThermoFisher |  | 27016021 |
| TMB | | KPL sureblue reserve |  | 5120-0081 |
| TWEEN |  | Sigma-Aldrich |  | P1379 |
| Monensin | | Biolegend |  | 420701 |
| Brefeldin A |  | Biolegend |  | 420601 |
| Zombie Aqua |  | Biolegend |  | 423102 |

| **Antibody** | **Clone** | **Source** | **Fluorochrome** | **Catalogue #** |
| --- | --- | --- | --- | --- |
| CD3 | 17A2 | Biolegend | Biotin  PB | 100244  100214 |
| CD11b | M1/70 | Biolegend | PE-Dazzle | 101256 |
| CD11c | N418 | Biolegend | APC-Cy7 | 117324 |
| CD16/32 | 2.4G2 | Biolegend | Purified | 101320 |
| CD19 | 6D5 | Biolegend | Biotin PB | 115504 115523 |
| CD45.1 | A20 | Biolegend | FITC AF700 | 110706 110724 |
| CD45.2 | 104 | Biolegend | AF700 | 109822 |
| CD102 | 3C4 | Biolegend | FITC | 105606 |
|  |  |  | AF647 | 105612 |
|  |  |  | Biotin | 105604 |
| CD209b | 22D1 | eBioscience/ThermoFisher | APC | 17-2093-82 |
| GATA6 | D61E4 | Cell Signalling Technologies | Purified | 5851S |
| F4/80 | BM8 | Biolegend | PE-Cy7  APC-Cy7 | 123114  123118 |
| FRβ | 10/FR2 | Biolegend | APC  PE | 153306  153303 |
| Sema4a | 5E3/SEMA4a | Biolegend | APC PE | 148406  148404 |
| CCR5 | HM-CCR5 | Biolegend | AF488 | 107008 |
| CD62L | MEL-14 | Invitrogen | SuperBright 702 FITC | 67-0621-82 11-0621-82 |
| MHC II (IA-IE) | M5/114.15.2 | Biolegend | AF700 | 107622 |
|  |  |  | PB APC-Cy7 | 107620  107628 |
| Ly6C | HK1.4 | Biolegend | BV711 | 128037 |
| SiglecF | ES22-10D8 | Miltenyi Biotec | Biotin | 130-101-861 |
| VSIG4 | NLA14 | eBioscience/ThermoFisher | PE-Cy7 | 25-5752-82 |
| Ly6G | 1A8 | Biolegend | Biotin PB | 127604 127612 |
| Tim4 | RMT4-54 | Biolegend | PE | 130006 |
|  |  |  | PE-Cy7 | 130010 |
|  |  |  | AF647 | 130008 |
| Streptavidin |  | Biolegend | BV650 | 405232 |
| Zenon anti-rabbit reagent |  | Molecular Probes | AF647 | Z25308 |
| Siglec F | E50-2440 | BD | BV421 | 562681 |
| Ki67 | REA183 | Miltenyi | FITC | 130-117-803 |
| CXCL13 | M1/70 | Invitrogen | APC | 17-7981-82 |
| CD11c  TNFα  Rat IgG1,Iso | N418  MP6-XT22  RTK2071 | Biolegend  Biolegend  Biolegend | APC-Cy7  BV421  BV421 | 117324 506328  400439 |
